# Proximity labeling reveals unique and shared interactomes of unmodified and pyroglutamate amyloid beta in human hippocampus in Alzheimer’s disease

**DOI:** 10.64898/2026.05.13.724866

**Authors:** Alia O. Alia, Kristy Urquhart, Hannah Carson, Bryan Killinger, Christopher Janson, Liudmila Romanova

## Abstract

Amyloid plaques are a hallmark neuropathological feature of Alzheimer’s disease (AD), composed of insoluble amyloid beta (Aβ) peptide. Aβ undergoes post-translational modifications that alter their biophysical properties, aggregation kinetics, and neurotoxicity, creating a heterogeneous pool of species that differentially affect AD pathogenesis. Pyroglutamate-modified Aβ (pEAβ) is a particularly aggregation-prone and proteolytically resistant variant that preferentially accumulates within plaque cores, is implicated in early plaque seeding, and is a major target of emerging anti-amyloid immunotherapies. However, the molecular environment surrounding pEAβ versus unmodified Aβ (pan-Aβ) in the human hippocampus remains incompletely defined. Here, we used Biotinylation by Antibody Recognition (BAR), an in-situ proximity labeling approach, to map and compare the protein-protein interactions (proteomes) of pEAβ and pan-Aβ in formalin-fixed postmortem human hippocampal tissue from pathologically confirmed AD cases and cognitively normal (CN) controls. Differential proteomic analysis identified 48 significantly enriched proteins in AD pEAβ captures, 28 in AD pan-Aβ captures, and 15 in CN pan-Aβ captures. Whereas no significant enrichment was detected in CN pEAβ captures, supporting pEAβ as a pathology-associated species. pEAβ in AD demonstrated the largest variant-specific signature with 31 unique proteins, pan-Aβ showed 11 unique proteins in AD, and 14 unique proteins in CN, 16 proteins were shared between AD pEAβ and AD pan-Aβ, with PCSK1N shared across AD pEAβ, and AD/CN pan-Aβ. Pathway enrichment analysis revealed broader biological disruptions linked to pEAβ, including synaptogenesis signaling, clathrin-mediated endocytosis, mitochondrial division signaling, and neurotransmitter release. Shared pathways included SNARE signaling, glutamatergic receptor signaling, and netrin signaling. These findings demonstrate that pEAβ engages an expanded, variant-specific interactome in human AD hippocampus and designate intracellular trafficking, synaptic signaling, and mitochondrial pathways as network-level vulnerabilities relevant to pEAβ pathology in AD. Notably, comparison of CN versus AD pan-Aβ further distinguished protein networks associated with physiological Aβ engagement versus pathological pan-Aβ deposition.

## Introduction

Alzheimer’s disease (AD) is a progressive neurodegenerative disorder and the most common cause of dementia [1]. A defining neuropathological hallmark of AD is the accumulation of amyloid plaques, which are primarily composed of sticky, insoluble aggregates of amyloid beta (Aβ) peptides [2]. These peptides cluster together, disrupting cell-to-cell communication and ultimately leading to synaptic loss and neuronal death [2]. Aβ peptides can aggregate into several structural forms, including oligomers, protofibrils, and fibrils that differ in biochemical characteristics, aggregation tendencies, and neurotoxic potential [2]. Fibrillar Aβ constitutes the main component of amyloid plaques, which is a major pathological hallmark of AD [3]. Aβ peptides are produced through the sequential cleavage of amyloid precursor protein (APP) by β-secretase and γ-secretase, generating various isoforms, including Aβ40 and the more aggregation-prone Aβ42 [2].

Despite ongoing debate regarding the causal role of Aβ accumulation in AD pathogenesis, Aβ remains a central biomarker and a principal therapeutic target [4]. Anti-amyloid immunotherapies have demonstrated the capacity to reduce cerebral Aβ burden [5]. Several monoclonal antibodies have been developed to target specific aggregation states or epitopes of Aβ. Aducanumab, a monoclonal antibody directed against aggregated Aβ, received accelerated approval from the U.S. Food and Drug Administration (FDA) based on its ability to reduce amyloid plaque load; however, clinical trials reported minimal cognitive benefit, and safety concerns were raised due to amyloid-related imaging abnormalities (ARIA) [6]. Similarly, Lecanemab, which preferentially binds Aβ protofibrils, has shown moderate efficacy in slowing cognitive decline but is also associated with ARIA-related adverse effects [6]. Therapeutically, the distinct composition of the Aβ pool has prompted the development of antibodies targeting specific Aβ variants. Donanemab, which selectively targets pyroglutamate-modified Aβ (pEAβ), a particularly pathogenic variant of Aβ, represents one of the most promising recent developments [7]. It has demonstrated robust plaque removal and significant reductions in cerebral amyloid burden in early-stage AD trials [8]. However, although donanemab effectively lowers amyloid load, its cognitive benefits have been modest, underscoring the need to better understand the mechanisms governing the formation, clearance, and metabolism of modified Aβ species [8]. These therapeutic outcomes demonstrate that although targeting Aβ modifies neuropathology, clinical benefits remain limited and often accompanied by significant risks. This highlights the unresolved question of which Aβ species and associated pathways critically drive disease progression and neurotoxicity, a major gap that continues to hinder the development of safe and effective treatments. Thus, advancing therapeutic success will require a more nuanced and mechanistic comprehension of

Aβ species dynamics within the complex molecular landscape of AD. Therefore, the goal of this study was to identify unique and overlapping molecular pathways involved in AD pathology of different Aβ species, unmodified Aβ and pEAβ.

As shown in Figure 1, pEAβ formation is facilitated by a set of enzymatic processes, most notably through the action of glutaminyl-peptide cyclotransferase (QPCT), which catalyzes the conversion of N-terminal glutamate (E) residues at position 3 or 11 of full-length Ab into pyroglutamate (pE)[9], [10]. This conversion occurs both intracellularly – primarily in acidic compartments such as the Golgi apparatus and presynaptic early and recycling endosomes – and extracellularly near amyloid plaques and synaptic clefts, where QPCT isoforms are also active [11], [12], [13].

**Figure 1.**
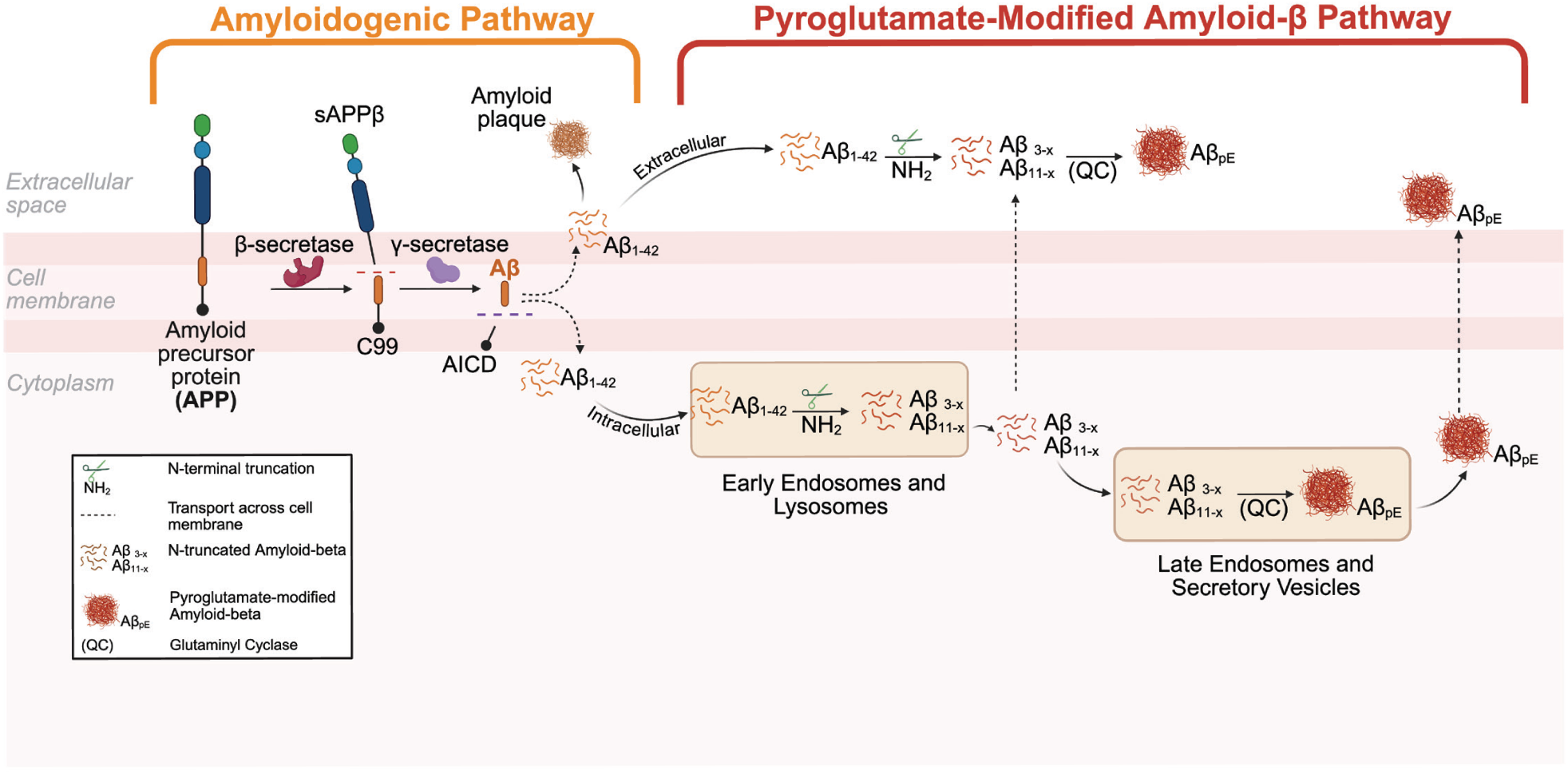
Cellular formation of pyroglutamate-modified amyloid beta (pEAβ). In the amyloidogenic pathway, amyloid precursor protein (APP) is sequentially cleaved by β-secretase (BACE1) and γ-secretase to generate amyloid beta (Aβ) peptides, primarily Aβ₁–₄₀ and Aβ₁–₄₂, which are secreted into the extracellular space where they aggregate into monomers, oligomers, fibrils, and plaques. In the pyroglutamate-modified Aβ pathway, Aβ peptides can be internalized into endosomal and lysosomal compartments, where they undergo further enzymatic processing by aminopeptidase A (APA), dipeptidyl peptidase 4 (DPP4), and neprilysin, yielding N-terminally truncated variants such as Aβ₃–₄₂ and Aβ₁₁–₄₂. These truncated peptides may then traffic to late endosomes or secretory vesicles, where the vesicular form of glutaminyl cyclase (QC) catalyzes the cyclization of N-terminal glutamate residues at positions 3 or 11 to form pEAβ, which is subsequently secreted into the extracellular space. Additionally, unmodified Aβ peptides released extracellularly can be cleaved at the N-terminus by BACE1, APA, DPP4, or meprin-β, producing truncated Aβ₃–₄₂ and Aβ₁₁–₄₂ species that are further converted to pEAβ through QC-mediated cyclization. Thus, pEAβ formation can occur both intracellularly and extracellularly, contributing to the overall amyloid burden and enhanced aggregation propensity characteristic of Alzheimer’s disease pathology.

The resulting cyclic pyroglutamate at the N-terminus blocks the free amine group, rendering pEAβ highly resistant to recognition and degradation by aminopeptidases and other Aβ-degrading enzymes such as insulin-degrading enzyme (IDE), neprilysin (NEP), endothelin-converting enzyme-1 (ECE-1), angiotensin-1 converting enzyme (ACE), and many other reported amyloid-degrading enzymes [14], [15]. As a result, pEAβ requires specialized degradation by pyroglutamyl peptidase (PGP), an enzyme typically restricted to intracellular and cytosolic compartments and largely absent in the extracellular brain environment where most plaque-associated pEAβ accumulates [16], [17]. This scarcity of PGP allows pEAβ to bypass physiological turnover and persist within brain tissue, ultimately contributing to plaque accumulation [17], [18], [19]. These biochemical properties render pEAβ not only more stable but also significantly more pathogenic than unmodified Aβ species, as it favors amyloidogenicity and resists proteolysis [16], [20], [21]. The chemical shift caused by pyroglutamate conversion of Aβ results in the loss of one positive charge and two negative charges. The net effect of these ionic changes dramatically increases the overall hydrophobicity of pEAβ, making it less soluble and more prone to self-associate in aqueous environments like the extracellular space of brain tissue [16], [21]. Numerous studies have shown that pEAβ aggregates faster than full-length Aβ, implicating the N-terminal truncation and pyroglutamation as key drivers of accelerated aggregation kinetics [22]. Consistent with its role in driving amyloid pathology, pEAβ is abundantly present in human AD tissue and is widely distributed across different types of Aβ deposits, including senile plaques, soluble oligomers, and other aggregate forms[23]. It preferentially accumulates in the core of amyloid plaques, standalone plaques, and intraneuronally in over half of AD cases, supporting its early role in plaque seeding [20], [24], [25], [26], [27]. *In vitro* studies further demonstrate that pEAβ exerts potent neurotoxic effects, inducing both apoptosis in neurons and necrosis in astrocytes at concentrations lower than those required for full-length Aβ to elicit similar toxicity [28]. Despite its pathogenic significance, the precise molecular mechanisms underlying pEAβ pathology remain poorly defined, largely due to the predominance of studies focusing on bulk-isolated aggregates or exclusively on the Aβ₁₋₄₂ isoform [13], [29]. Such approaches obscure both the biochemical diversity of Aβ variants and the spatial and molecular context in which these peptides accumulate, limiting our understanding of the distinct microenvironments that influence their pathogenic behavior.

Together, these observations underscore the need to resolve the spatial and molecular diversity of amyloid species within the human brain. Multiple proteomic studies of bulk-isolated plaques or homogenized brain tissue have been conducted to identify pathogenic pathways and proteins involved in amyloid pathology development[30], [31]. These approaches have revealed variations in amyloid plaque composition and disruptions in protein networks associated with synaptic, mitochondrial, and immune dysfunction [31], [32], [33]. However, these studies have largely overlooked post-translationally modified Aβ forms and the spatial relationships among distinct Aβ species, as well as the distinction between functional interactions and nonspecific associations. In this study, we addressed this gap by comparing the protein–protein interaction networks (interactomes) of unmodified Aβ and pyroglutamate-modified Aβ, a therapeutically relevant amyloid species. We employed a spatial proteomic strategy, Biotinylation by Antibody Recognition (BAR), which enables *in situ* labeling of proteins in close proximity to a target antigen within native tissue environments. This method preserves insoluble and membrane-associated components, allowing the capture of physiologically relevant protein interactions. Given that pEAβ and Aβ pathologies only partially overlap, engaging distinct molecular pathways, such spatially resolved analysis is critical. Our findings demonstrated that pEAβ and unmodified Aβ species are associated with distinct molecular pathways in human brain, revealing variant-specific protein interactions that may differentially contribute to disease progression. In addition, we identified a subset of interacting proteins shared between pEAβ and Aβ, suggesting convergence of certain physiological mechanisms underlying amyloid pathology formation.

## Methods

### Human brain tissue selection and preparation

Specimens were obtained from the Banner Sun Health Research Institute brain bank from the Arizona Study of Aging and Neurodegenerative Disease (AZSAND). Donors enrolled in the study underwent comprehensive clinical assessments during life, which included neurological exams, neuropsychological tests, and medical record reviews. Upon death, the autopsy protocol is initiated with a median post-mortem interval (PMI) of 3 hours to optimize tissue preservation [34]. Brain extraction was performed using an oscillating electric saw to open the calvarium and expose the intact brain, followed by the extraction of the brain with the olfactory bulbs and pituitary gland still intact. Gross pathological features are then documented and photographed. The brain is then immediately divided so that the cerebellum and brainstem are separated from the cerebrum, and the cerebrum is coronally sectioned into 1cm slices using a custom slicing jig. The hippocampus is systemically sampled at multiple levels (head and body) and evaluated histologically using standardized criteria, including CERAD (plaques), Braak (neurofibrillary tangles), and Thal (amyloid distribution) staging, immunohistochemistry for phosphorylated a-synuclein for Lewy pathology, and TDP-43 for frontotemporal lobar degeneration. The hippocampus from the left hemisphere was immersion fixed in 4% formaldehyde for 48 hours at 4°C, followed by a thorough rinse in 0.1M phosphate buffer for 24 hours to remove excess fixative. The tissue was then cryoprotected in a solution containing 20% glycerol and 2% DMSO for 48 hours at 4°C. The tissue was then embedded in OCT and frozen in dry ice-cooled isopentane and stored at -80°C until processing. Next, using a freezing microtome, 40-80 µm thick sections were sliced and collected in cryoprotectant solution for long-term storage at -20°C. [34] Human hippocampus tissue from five AD confirmed cases, as well as five cognitively normal cases, was selected. Neuropathological classification followed standard CERAD and Braak criteria. All AD samples met criteria for definite AD based on CERAD NP scoring and exhibited “frequent” neuritic plaques, Braak stage V-VI or higher, and evidence of cerebral amyloid angiopathy (CAA). CN samples were limited to cases with Braak stages I-III and no reported CAA. The hippocampus was selected because it represents a well-defined medial temporal lobe structure that is highly relevant to AD and is routinely examined in postmortem AD studies[35]. In addition to its biological relevance, the hippocampus offers practical advantages for comparative analysis because its anatomy is relatively well described, whereas cortical Aβ burden is more regionally heterogeneous, and selection of a single cortical section can be difficult to standardize across patients [35], [36]. For these reasons, focusing on the hippocampus provides a biologically meaningful and methodologically consistent approach for studying Aβ-associated pathology in human tissue.

### Immunohistochemistry (IHC) and Imaging

Free-floating formalin-fixed hippocampal sections was washed three times in dilution media (DM; 150 mM NaCl, 8 mM Tris base, 41 mM Tris-HCl, and 0.05% Triton X-100) followed by antigen retrieval consisting of an incubation in heated sodium citrate buffer for 30 minutes in a water bath. First, the tissues were washed in room temperature sodium citrate buffer for five minutes, then washed in a 95°C heated bath for 30 minutes. Next, the samples were placed on ice for 15 minutes, followed by two washes in DM. Endogenous peroxidase inhibition and serum blocking were performed, consisting of DM, 3% goat serum, 2% BSA, and 0.4% Triton X-100 for 1 hour. Primary antibody, rabbit monoclonal to β-amyloid pE3 peptide (D5N5H, Cell Signaling catalog # 14975, dilution 1:1000) was prepared similarly to blocking with hydrogen peroxide and 10% sodium azide and added to tissue, followed by overnight incubation at 4°C. Tissue was washed 3 times in DM, followed by goat anti-rabbit HRP-secondary antibody incubation at 1:1000 (ThermoFisher Scientific catalog # 31460) for 1 hour. Tissue was then incubated in 0.05M borate buffer for 10 minutes, followed by secondary antibody incubation containing 0.003% hydrogen peroxide and Biotium CF568 Tyramide (Biotium catalog # 92173, dilution:1:2000) for 30 minutes. All steps were repeated for a second round of antibody staining in anti-β-Amyloid, 1-16; 6E10 (BioLegend catalog # 803014, dilution 1:10,000), followed by goat anti-mouse HRP-secondary antibody incubation at 1:1000 (ThermoFisher Scientific catalog #31430) for 1 hour, washed, and incubated in AlexaFluor 647 goat anti-mouse (ThermoFisher Scientific catalog # A-21235, dilution 1:2000) for 1 hour. The tissue was then mounted to slides using anti-fade Prolong Gold media (ThermoFisher Scientific catalog # p36984) and imaged on a Nikon A1R confocal microscope. Images were acquired using Nikon CFI60 Plan Fluor objectives for 20x and 40x (air) magnifications, and a CFI60 Plan Apochromat objective for 60x oil immersion imaging.

### IHC with DAB Detection

Free-floating formalin-fixed hippocampal sections from AD and CN tissue were processed for chromogenic IHC using 3,3’-diaminobenzidine (DAB) detection, following the same fixation, blocking, and washing steps described above for fluorescence IHC. Antigen retrieval was performed in 10mM sodium citrate buffer (pH 6.0) containing 0.05% Tween-20 at 90-95°°C for 10 min, followed by cooling on ice and 3 washes in DM. Endogenous peroxidase activity was quenched for 30 minutes using 0.3% hydrogen peroxide and 0.1% sodium azide in DM. Sections were then incubated overnight at 4°°C with the primary antibody β-amyloid pE3 peptide (1:1000) or 6E10 (1:10,000) diluted in blocking buffer. Goat anti-rabbit and goat anti-mouse HRP-conjugated secondary antibodies (1:200) were prepared in blocking buffer, and the sections were left to incubate for 1 hour. For chromogenic development, sections were equilibrated in imidazole–acetate buffer containing sodium acetate trihydrate (Fisher Scientific catalog # S209-500), imidazole (Millipore Sigma catalog # I2399), and milli-Q water; adjusted pH to 7.2-7.4 with glacial acetic acid. DAB reactions were performed in unp-H’ed imidazole–acetate buffer containing 3,3’-diaminobenzidine tetrahydrochloride hydrate (Millipore Sigma catalog # D5637) and 1% hydrogen peroxide. For nickel enhancement, nickel (II) sulfate hexahydrate (Millipore Sigma catalog # N4882) was added. The reaction was monitored visually and terminated after 10 minutes by rinsing in pH-adjusted imidazole–acetate buffer, followed by 2 washes in DM and 2 washes in PBS. Sections were mounted onto glass slides using Cytoseal 60 mounting medium (Fisher Scientific catalog # 23-244257). Images were acquired in brightfield using 20x magnification.

### Hematoxylin and Eosin (H&E) Staining

Free-floating formalin-fixed hippocampal sections were processed for H&E staining following standard histological procedures. Briefly, free-floating sections were mounted onto Superfrost Plus microscope slides (Fisher Scientific catalog # 22-034979), air dried for 1 hour, and hydrated in tap water for 4 minutes. Slides were then stained in hematoxylin (Fisher Scientific catalog # 22-050-203) for 4 minutes, differentiated by three dips in 1-2% acetic acid prepared in 70% ethanol, and rinsed gently under running tap water for 3 minutes. Slides were immersed in Scott’s special tap water concentrate (Leica Biosystems catalog # 3802900) for 2 minutes to blue nuclei, followed by an additional 2-minute tap water rinse. Sections were counterstained in eosin (Azer Scientific catalog # ES737) for 2 minutes and then dehydrated through graded ethanols (70%, 95%, 100%) for 2 minutes each. Slides were cleared in two subsequent xylene baths (Fisher Scientific catalog # X3P-1GAL), 5 minutes each under a chemical fume hood. Slides were then coverslipped with Cytoseal 60 mounting medium (Fisher Scientific catalog # 23-244257). All slides were allowed to dry overnight under a chemical hood before imaging. Images were acquired in brightfield using CFI60 Plan Fluor in 4x magnification.

### Biotinylation by Antibody Recognition (BAR) Labeling of Human Tissues

Free-floating formalin-fixed hippocampal sections were washed three times in dilution media (DM), followed by antigen retrieval consisting of an incubation in heated sodium citrate for 30 minutes in a water bath. First, the tissues were washed in room temperature sodium citrate buffer for five minutes, then washed in a 95°C heated bath for 30 minutes. Next, the samples were placed on ice for 15 minutes, followed by two washes in DM. Endogenous peroxidase inhibition and serum blocking were performed for 1 hour. This was performed in DM, goat serum, bovine serum albumin, and Triton X-100. Antibody selection was an important consideration in this study. Clone 6E10 was selected to detect N-terminal Aβ-containing material as it recognizes an epitope within Aβ residues 3-8. In parallel, the pyroglutamate Aβ antibody was selected to detect pE3-Aβ, with manufacturer validation indicating it does not cross-react with non-pyroglutamate Aβ forms[37]. Primary antibody solutions were prepared for each pulldown condition in blocking solution (β-amyloid pE3 peptide Rb 1:1000, 6E10 Ms 1:10,000) with hydrogen peroxide and 10% sodium azide and left to incubate overnight at 4°C. Tissue was washed 3 times in DM followed by HRP-secondary antibody incubation for 1 hour in either goat anti-rabbit HRP (Invitrogen catalog # 31462) or goat anti-mouse HRP (Invitrogen catalog # G21040), followed by two washes in DM. Tissues were then incubated for 10 minutes in 0.05M borate buffer. Sections were then washed in DM 3 times for 10 minutes and transferred to 1.5mL Eppendorf tubes, followed by quick centrifugation, and any residual buffer was removed. 0.5mL reversal buffer (SDS, 0.5M EDTA, Tris-base, and NaCl) was added to each tube, and samples were briefly sonicated on low power to disperse tissue. 5uL of 100 uM PMSF was added to the samples, followed by heating at 98°C for 30 minutes. Samples were removed from heat and allowed to cool, then vortexed, followed by centrifugation at 21,120 xg for 20 minutes at room temperature. Extracted proteins were collected into 15mL conical and 4.5 mL TBST (1XTBS, 1% Triton X-100 in milli-Q water) was added. Pierce Streptavidin Magnetic Beads (ThermoFisher Scientific catalog # 88817) were prepared by surface washing in milli-Q water and resuspension in 100 μL TBST. Samples were vortexed, followed by the addition of 40uL prepared magnetic streptavidin beads. Samples were placed on the tube rotator and rotated for 1 hour at room temperature. Tubes were placed on a magnetic stand for 2 minutes to allow the beads to be drawn out of solution, followed by the removal of the buffer. 10mL of wash buffer (1XTBS, 1% Triton X-100, SDS, EDTA in milli-Q water) was added to each tube, followed by 30 minutes of rotation. This step was repeated two more times, followed by rotation in a 10mL wash buffer overnight. The following day, tubes were placed on a magnetic stand for 2 minutes, followed by the removal of the buffer. Beads were resuspended in 1mL wash buffer in a 1.5mL tube, followed by two surface washes in milli-Q water. The beads were spun down, and all liquid was drawn out. 80uL of 1X SDS-PAGE buffer containing 4X Bolt LDS Sample buffer (ThermoFisher Scientific catalog # B0007), 10X Bolt Sample Reducing Agent (ThermoFisher Scientific catalog # B0009) and milli-Q water was added to each sample. The samples were vortexed, spun down, and placed on a heat block set to 98°C for 10 minutes. Samples were removed, vortexed, and spun down, then placed on a magnetic stand for 2 minutes. The eluent from each tube was collected and placed into low-bind tubes, and the beads were saved.

### SDS Electrophoresis

Samples were subjected to gel electrophoresis to prepare proteins for mass spectrometry in-gel digestion. Bolt Bis-Tris 4-12% gels (ThermoFisher Scientific catalog # NW04125BOX) were prepared in 1X MOPS running buffer. 5uL of 1X SDS-PAGE buffer standard was loaded into the first well, followed by 40uL of each sample into the remaining wells while skipping a well in between samples to avoid cross-mixing. Empty wells were loaded with 40 μL 1X loading buffer. Gels were run at 200V for approximately 5 minutes or until the sample had completely entered the gel. Gels were removed and fixed for 1 hour at room temperature in fixation solution (50% ethanol, 10% acetic acid). Gels were then washed in several changes of milli-Q water until the gels had swollen back to their original size. 100mL colloidal SimplyBlue SafeStain (ThermoFisher Scientific catalog # LC6060) was added to each gel, and gels were covered loosely and heated in a microwave until the solution began to boil. Gels were incubated in a heated colloidal Coomassie blue stain for 8 minutes. Gels were removed from the stain solution and washed several times with milli-Q water until the backgrounds ran clear. Gels were placed on a clear acrylic surface, and samples were excised using a razor. Samples were placed into clean 1.5mL tubes containing 500uL milli-Q water and stored at 4°C until submitted for mass spectrometry analysis.

### Dot blot

Biotin Dot blots were performed to estimate target enrichment. PVDF membranes were activated in 100% methanol for 30s followed by equilibration in milli-Q water. Membranes were patted with Kim wipes to remove any extra moisture, and a spot of 1uL of each sample was loaded onto the membrane. Membranes were allowed to dry completely, followed by reactivation in 100% methanol for 30s and equilibration in milli-Q water. Membranes were blocked in blocking buffer containing 5% BSA in wash (1XTBS, 0.1% Tween-20) for 1 hour. An ABC complex solution was prepared in the wash buffer, and membranes were incubated in the solution for 30 minutes. Membranes were washed in wash buffer 3 times, followed by development in Pierce ECL Western Blotting Substrate (ThermoFisher Scientific catalog # 32106) and imaged on a ChemiDoc imager (BioRad catalog # 12003153). Antibody detection blots were performed to confirm antibody specificity and were prepared similarly to biotin dot blots, followed by 3 washes in wash buffer. Membranes were then incubated in primary antibody, either containing β-amyloid pE3 peptide 1:600 or 6E10 1:3,350 in blocking solution overnight at 4°C. The following day, membranes were washed 3 times in wash buffer followed by incubation in secondary antibody goat anti-rabbit HRP 1:10,000 or goat anti-mouse HRP 1:5000 in blocking buffer for 1hour. Membranes were then washed 3 times in wash buffer, followed by development in SuperSignal West Atto Ultimate Sensitivity Substrate (ThermoFisher Scientific catalog # A38555) and imaged on a ChemiDoc imager.

### Mass spectrometry

Proteins were excised from polyacrylamide gels and subjected to in-gel digestion for subsequent mass spectrometry analysis. The gel pieces were initially destained by washing with 100 mM ammonium bicarbonate (AmBic) and 50% acetonitrile (ACN) to remove background staining, followed by a dehydration step with 100% ACN. The gel was then reduced with 10 mM dithiothreitol (DTT) in 100 mM AmBic and alkylated with 100 mM iodoacetamide (IAA) in 100 mM AmBic to break disulfide bonds and prevent reformation. After thorough washing, trypsin was added at a 1:50 ratio (enzyme: substrate) and incubated overnight at 37°C for protein digestion into peptides. The peptides were extracted in a series of solvent washes, beginning with 50% ACN/5% formic acid, followed by 80% ACN/5% formic acid, and finally 100% ACN. The resulting supernatants were pooled, concentrated using a speed vacuum, and resuspended in 5% acetonitrile and 0.1% Formic acid. Peptide analysis was performed using a Vanquish Neo nano-LC system coupled to an Orbitrap Exploris 240 hybrid quadrupole-Orbitrap mass spectrometer (Thermo Fisher Scientific). The peptides were separated on a UHPLC C18 column (Ion Opticks, AUR3-15075C18-CSI), with 1 µg of peptide sample injected per analysis. Ionization was achieved through electrospray using a Nanospray Flex Ion Source (Thermo Fisher, ES071) set to a positive spray voltage of 2.2 kV. Peptides were eluted at a flow rate of 200 nL/min using an increasing organic gradient to separate them based on hydrophobicity. The mobile phase consisted of buffer A (0.1% formic acid in LC-MS grade water) and buffer B (80% acetonitrile, 19.9% LC-MS grade water, 0.1% formic acid). The total run time was 120 minutes. The Orbitrap mass spectrometer was operated in positive polarity using Xcalibur software. Full scan (MS1) settings included a mass range of 350-1600 m/z, an RF lens voltage of 60%, Orbitrap resolution at 120,000, a normalized AGC target of 300%, and a maximum injection time of 25 ms, with a 5E3 intensity threshold. Data-dependent acquisition (DDA) was carried out using TopN with higher-energy collisional dissociation (HCD) for precursor ions with charge states between 2+ and 5+. MS2 parameters were dynamic exclusion for 30 seconds, a mass tolerance of 5 ppm, a 2-second cycle time, a 1.5 m/z isolation window, 30% normalized collision energy, Orbitrap resolution of 15,000, a normalized AGC target of 100%, and a maximum injection time of 50 ms.

### Data and Proteomic Analysis

Mass spectrometry files (.raw) were converted to Mascot generic format (.mgf) using the Scripps RawConverter program and then analyzed with the Mascot search engine (Matrix Science, version 2.5.1). MS/MS spectra were queried against the *Homo sapiens* SwissProt database. Search parameters included cysteine carbamidomethylation as a fixed modification, and variable modifications for methionine oxidation, deamidation of asparagine and aspartic acid, and acetylation of protein N-termini. Label-free proteomics experiment data were analyzed using Scaffold (version 5.3.3). A protein and peptide threshold of 5% false discovery rate (FDR) was used, and a minimum number of 2 peptides per protein was required for quantification. The normalized total ion current (TIC) for all detected peptides in all samples was exported as an Excel table. The resulting table of TIC values was analyzed in R (version 4.2.2). Contaminant peptides with an accession ID starting with “Cont_” as well as keratin were removed from the table. Decoy peptides with accession ID ending in “-DECOY” were also removed. The resulting filtered peptide table was further filtered by keeping those peptides that had less than or equal to 70% samples with no signal within each primary antibody pulldown group. Specifically, peptides were retained if they exhibited a detectable signal in 3 or more of the 10 samples for the pEAβ or pan-Aβ groups and a detectable signal in 6 or more of the 20 samples from the no-primary antibody groups. The final dataset represented the union of peptide features retained across all groups. Peptide-level total ion current (TIC) values were then log_2_ transformed using a pseudo-count of 1 to stabilize variance and normalize data distribution. Differential analysis between groups was quantified in R (version 4.2.2) using the limma package under a general linear model (GLM) approach [38]. Analyses were carried out comparing the primary antibody pulldown-groups (pEAβ, pan-Aβ, and none), the pathology (AD and CN), and their interaction. Pairwise comparisons were also performed between all combinations of antibody and pathology groups to identify variant and disease-specific protein changes. Multiple testing correction was applied using the Benjamini-Hochberg FDR procedure, and proteins were considered significantly differentially expressed at an adjusted p-value (FDR) [39]. Data visualization and figure generation were performed in R (version 4.2.2) and GraphPad Prism (version 10.2.0). Protein interaction networks were generated using the STRING database (version 12.0) and visualized in Cytoscape (version 3.10.0).

## Results

### Sample selection for BAR proximity labeling in human hippocampus

We analyzed postmortem hippocampal tissue from 10 individuals obtained through the Arizona Study of Aging and Neurodegenerative Diseases (AZSAND) brain bank. The cohort included five cognitively normal (CN) cases and five pathologically confirmed AD cases (Table 1). Among the CN group, donors ranged from 71-90 years of age (4 males, 1 female) with a median postmortem interval (PMI) of 3.8 hours, Braak stages I-III, sparse to absent amyloid plaques, and no evidence of cerebral amyloid angiopathy (CAA). The AD group included donors aged 78-89 years (3 males, 2 females) with a median PMI of 3.1 hours, frequent neuritic plaques, Braak stages V-VI, and confirmed CAA. The inclusion of CN cases with low amyloid burden was critical for investigating non-pathological interactions of Aβ species in cognitively normal tissue, allowing for comparison with pathological interactions surrounding Aβ species in AD tissue. The relatively short PMI (2-4 hours across all samples) ensured optimal tissue preservation for downstream proteomic analyses.

**Table 1.** Demographic and neuropathological characteristics of human hippocampus specimens. Summary of donor sex, age at death, PMI, neuropathological diagnoses, CAA status, total and density of neuritic plaques, and Braak stage for all CN and AD specimens used in this study. Ages ranged from 71 to 90 years of age, with four CN males, 3 AD males, 1 CN female, and 2 AD females. The average PMI for all specimens was approximately 3.5 hours, consistent with rapid autopsy protocols. CAA was present in all AD samples and absent in all CN samples. All CN samples exhibited low plaque burden (sparse to zero) and Braak stages I-III, while AD cases exhibited significant plaque burden (frequent) and Braak stages V-VI.

| Case ID | Sex | Age at death | PMI | Status | CAA | Plaque total and density | Braak |
| --- | --- | --- | --- | --- | --- | --- | --- |
| 17-50 | M | 90 | 3.41 | CN | No | 0.5 (Sparse) | I |
| 18-62 | M | 80 | 4.08 | CN | No | 0 | II |
| 18-78 | M | 71 | 3.5 | CN | No | 0 | III |
| 19-48 | M | 71 | 4.05 | CN | No | 0 | II |
| 22-16 | F | 87 | 4.03 | CN | No | 0 | III |
| 20-10 | M | 78 | 2.28 | AD | Yes | 14 (Frequent) | V |
| 20-25 | M | 89 | 4.17 | AD | Yes | 15 (Frequent) | VI |
| 20-66 | F | 85 | 2.87 | AD | Yes | 14 (Frequent) | V |
| 21-43 | F | 86 | 3 | AD | Yes | 15 (Frequent) | V |
| 21-46 | M | 87 | 3.62 | AD | Yes | 14 (Frequent) | V |

### Experimental group design for BAR proximity labeling in human hippocampus

For each pathological condition (CN and AD), we generated four groups based on the targeted Aβ variant and the inclusion of technical controls (Table 2).

**Table 2.** Allocation of samples to labeling conditions and control groups. Summary of experimental group design for BAR proximity labeling in human hippocampal tissue. Each pathological condition (AD and CN) included two primary proximity labeling groups, targeting pEAβ or pan-Aβ using the respective antibody (pE3 peptide or 6E10): pEAβ (AD), pan-Aβ (AD), pEAβ negative (CN), and pan-Aβ negative (CN). Corresponding no primary (no antibody) controls were included for each labeling condition to enable background correction and exclude non-specific interactions: pEAβ no primary (AD), pan-Aβ no primary (AD), pEAβ negative no primary (CN), and pan-Aβ negative no primary (CN).

| Group | Description |
| --- | --- |
| pEAβ | AD, proximity labeling using pE3 peptide |
| pan-Aβ | AD, proximity labeling using 6E10 |
| pEAβ no primary | AD, no primary antibody; background correction for pEAβ group |
| pan-Aβ no primary | AD, no primary antibody; background correction for pan-Aβ group |
| pEAβ negative | CN, proximity labeling using pE3 peptide |
| pan-Aβ negative | CN, proximity labeling using 6E10 |
| pEAβ negative no primary | CN, no primary antibody; background correction for pEAβ negative group |
| pan-Aβ negative no primary | CN, no primary antibody; background correction for pan-Aβ negative group |

Samples were processed using BAR to achieve proximity labeling of protein-protein interactions associated with either pEAβ or unmodified pan-Aβ species. Within the AD condition, two primary labeling groups were established (pEAβ and pan-Aβ), each corresponding to proximity labeling with the respective antibody in AD tissue that is abundant in pathological Aβ species. To account for non-specific protein labeling or any activity unrelated to target proximity, “no primary” control groups were included in parallel to pEAβ and pan-Aβ, identified as pEAβ no primary and pan-Aβ no primary. Data from the matched no primary groups were used for background subtraction, allowing the exclusion of non-specific signals and retention of true proximity-based interactors. An equivalent design was applied to the CN specimens, generating the groups pEAβ negative and pan-Aβ negative that correspond to proximity labeling with the respective antibody in CN tissue that contains sparse to zero Aβ species. CN no primary groups were also included to account for non-specific protein labeling and unrelated target proximity activity (pEAβ negative no-primary and pan-Aβ negative no-primary). Similarly, data from the matched no-primary controls were used for background subtraction to ensure the final dataset is reflective of true proximity-based interactors. Although CN tissue exhibits minimal to no Aβ pathology, any interactions exhibited in this tissue may provide valuable context for understanding baseline, non-pathological Aβ protein interactions. Comparing CN and AD interactomes enabled the distinction between baseline Aβ-associated pathways and those specifically enriched in the diseased state, thereby contributing to the clarification of molecular mechanisms that may transition from normal Aβ turnover to pathological aggregation.

### Aβ interactome in human hippocampus

To investigate the proteomic landscape in the formation of plaques containing unmodified and pEAβ species in human hippocampus, we applied biotinylation by antibody recognition (Fig. 2A). This proximity labeling method enables in situ biotinylation of proteins located in proximity of a specific antigen in fixed tissue, preserving native structural and biochemical context. The workflow involves antigen retrieval, antibody-guided horseradish peroxidase (HRP) labeling, and biotin-tyramide deposition, capturing proteins spatially associated with the target antigen.

**Figure 2.**
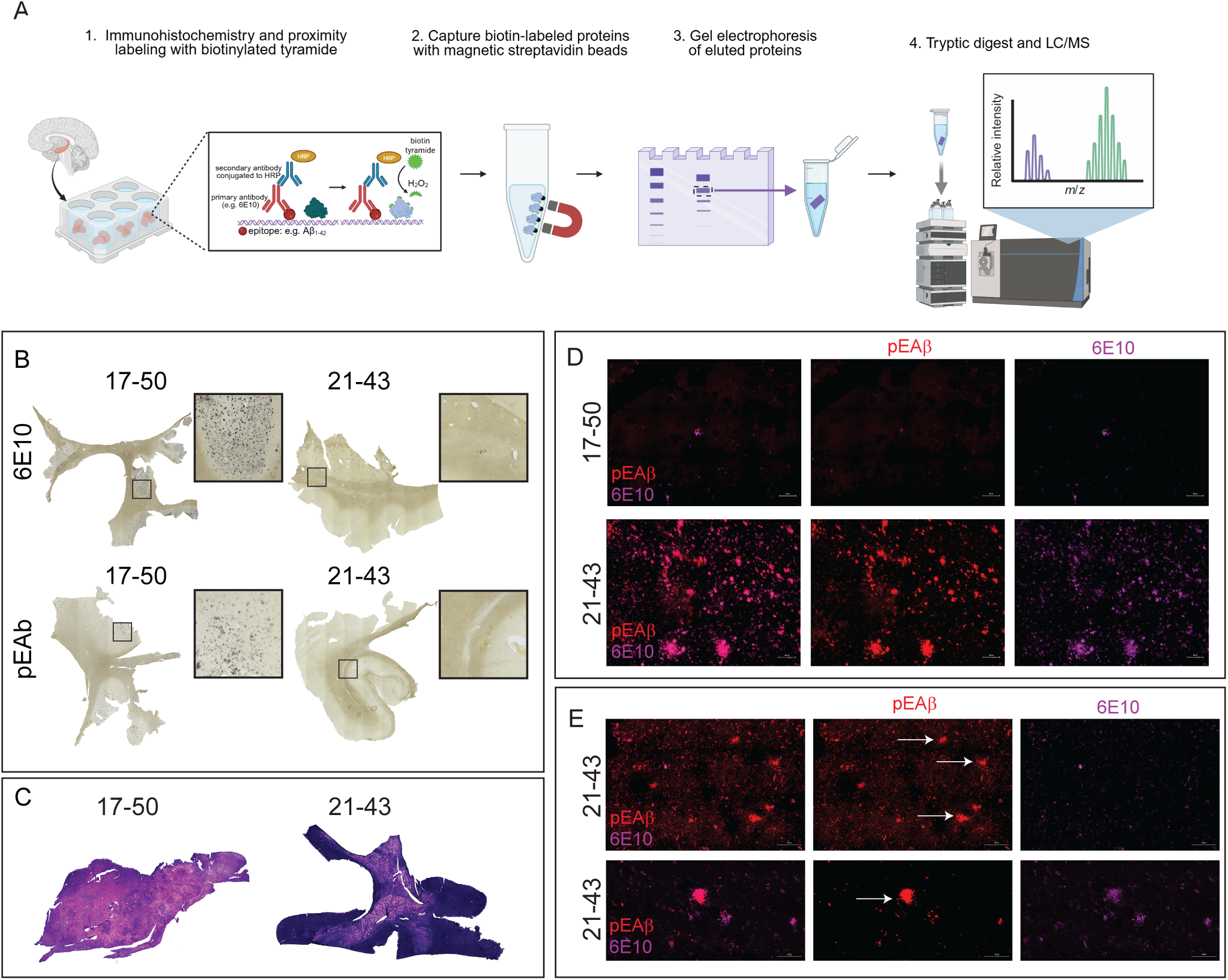
BAR-mediated capture and distribution of Aβ and pEAβ plaques in human hippocampal specimens. (A) Overview of the BAR workflow. Fixed floating sections of human hippocampus were incubated with anti-pEAβ and anti-pan-Aβ (6E10) antibodies. Secondary antibodies conjugated to horseradish peroxidase (HRP), in the presence of biotinyl-tyramine, catalyzed the labeling of proteins in proximity to the antibody–Aβ complexes. Following reversal of cross-linking, biotin-labeled Aβ-associated protein complexes were purified using streptavidin magnetic beads, separated by electrophoresis, excised, and analyzed via LC–MS. (B) Immunohistochemical staining of hippocampal sections from AD case (21–43) and a cognitively normal (CN) control (17–50) showing characteristic immunoreactivity with 6E10 and pEAβ antibodies. Abundant Aβ and pEAβ plaques are evident in AD specimens, whereas only sparse plaques are observed in CN tissue, although some residual plaque pathology remains. (C) H&E staining of AD and CN hippocampal sections showing preserved overall tissue morphology. (D–E) Immunohistochemical labeling of CN (17–50) and AD (21–43) hippocampal sections highlighting the differential deposition of pEAβ (red) and pan-Aβ (purple) at 20× (D) and 60× (E) magnification. Both pEAβ and pan-Aβ form distinct as well as mixed plaques (arrows), reflecting their coexistence and overlapping distribution within hippocampal tissue. Scale bar = 100um.

For selective labeling of pEAβ, we used the rabbit monoclonal antibody β-Amyloid (pE3 Peptide) (D5N5H), which exhibits high specificity for the N-terminal pyroglutamate residue (position 3) of Aβ and does not cross-react with unmodified or other truncated forms of Aβ peptide [37], [40]. To label unmodified Aβ, we employed the mouse monoclonal antibody 6E10, which recognized residues 1-16 of the Aβ sequence, which is an epitope retained in the full-length peptide [41], [42]. This antibody design allowed us to distinguish variant-specific protein environments and compare the interactomes of distinct Aβ species.

Before BAR labeling, tissues were characterized by IHC using the same antibodies to confirm that each specimen reflected its assigned pathological group and to evaluate the distribution and organization of Aβ deposits (Fig. 2 B-E). In AD specimens, dense plaques were abundant throughout the hippocampus with both diffuse and compact morphologies (Fig. 2B). pEAβ immunoreactivity was prominent in plaque cores, whereas pan-Aβ immunostaining outlined diffuse borders. Across AD cases, plaque density and morphology were comparable. In contrast, CN hippocampal tissue exhibited sparse, small, and isolated pan-Aβ deposits that lacked dense core morphology. pEAβ staining in CN tissue was minimal to a single specimen (17-50) and absent in all other specimens; this is consistent with low amyloid burden and the absence of mature Aβ plaques (Fig. 2B). To further assess the spatial relationship between pEAβ and unmodified Aβ species, we performed dual immunofluorescence staining on adjacent hippocampal sections using both antibodies. AD tissues displayed a heterogeneous mixture of pEAβ and unmodified Aβ plaques (Fig. 2D-E). We also observed pEAβ dominant plaques that exist on their own. CN tissues displayed minimal unmodified Aβ plaque deposition with no observed pEAβ deposits. Together, these observations indicate that pEAβ and unmodified Aβ exhibit overlapping yet distinct spatial distributions, supporting the presence of structurally and biochemically distinct Aβ species within hippocampal plaques, reinforcing the variant-specific nature of Aβ aggregation in AD.

### Mass spectrometry analysis of pEAβ and unmodified Aβ BAR capture revealed distinct and unique interactomes in human hippocampus

Immunohistochemical analysis confirmed the presence of the target pathology in the tissues. The same tissues were used for proximity labeling. The labeling was performed to capture proteins spatially associated with each Aβ species as previously described [43], [44]. All AD samples and CN samples were processed in parallel with matched “no primary” controls for the exclusion of non-specific protein interactors during downstream analyses. Prior to mass spectrometry, we conducted a series of quality control (QC) assays to evaluate the labeling efficiency and protein recovery. First, specimens are run on Coomassie-stained SDS-PAGE gels to estimate the relative protein concentration of each capture. Samples were run alongside BSA standards, enabling visual estimation of total protein yield and identification of samples with insufficient recovery. QC images were provided to the mass spectrometry core to guide the appropriate digestion volumes. Next, biotin dot blots were performed to verify successful proximity labeling and ensure robust enrichment of biotinylated proteins. Each sample was spotted onto PVDF membranes, probed with an avidin-HRP/ABC complex, and developed by chemiluminescence (Additional file 1A). All AD and CN samples produced strong biotin signals relative to their paired no-primary controls, confirming efficient labeling of nearby proteins. Finally, antibody target dot blots were used to validate primary antibody specificity and confirm that labeling was restricted to the intended Aβ species (Additional file 1B).

Following mass spectrometry, raw peptide tables from all capture groups (pEAβ, pan-Aβ, pEAβ negative, and pan-Aβ negative) underwent background correction using their matched no primary controls. This entailed the removal of proteins that appeared in the in the no primary controls for the corresponding capture group to ensure the retention of true labeling events rather than non-specific interactions. After QC and background subtraction, a total of 557 proteins were identified across all human AD and cognitively normal samples (Additional file 2). We found 48 significantly enriched proteins in AD samples with pEAβ antibody capture, 28 significantly enriched in AD samples within pan-Aβ AD, and 15 significantly enriched proteins within pan-Aβ CN tissues (Fig. 3A and Table 3). No proteins were significantly enriched within pEAβ in cognitively normal samples. Among the significantly enriched proteins, unique interactors were detected across capture conditions, with 31 uniquely enriched proteins in AD pEAβ, 11 proteins unique to AD pan-Aβ, and 14 uniquely enriched proteins in CN pan-Aβ specimens (Fig. 3B).

**Figure 3.**
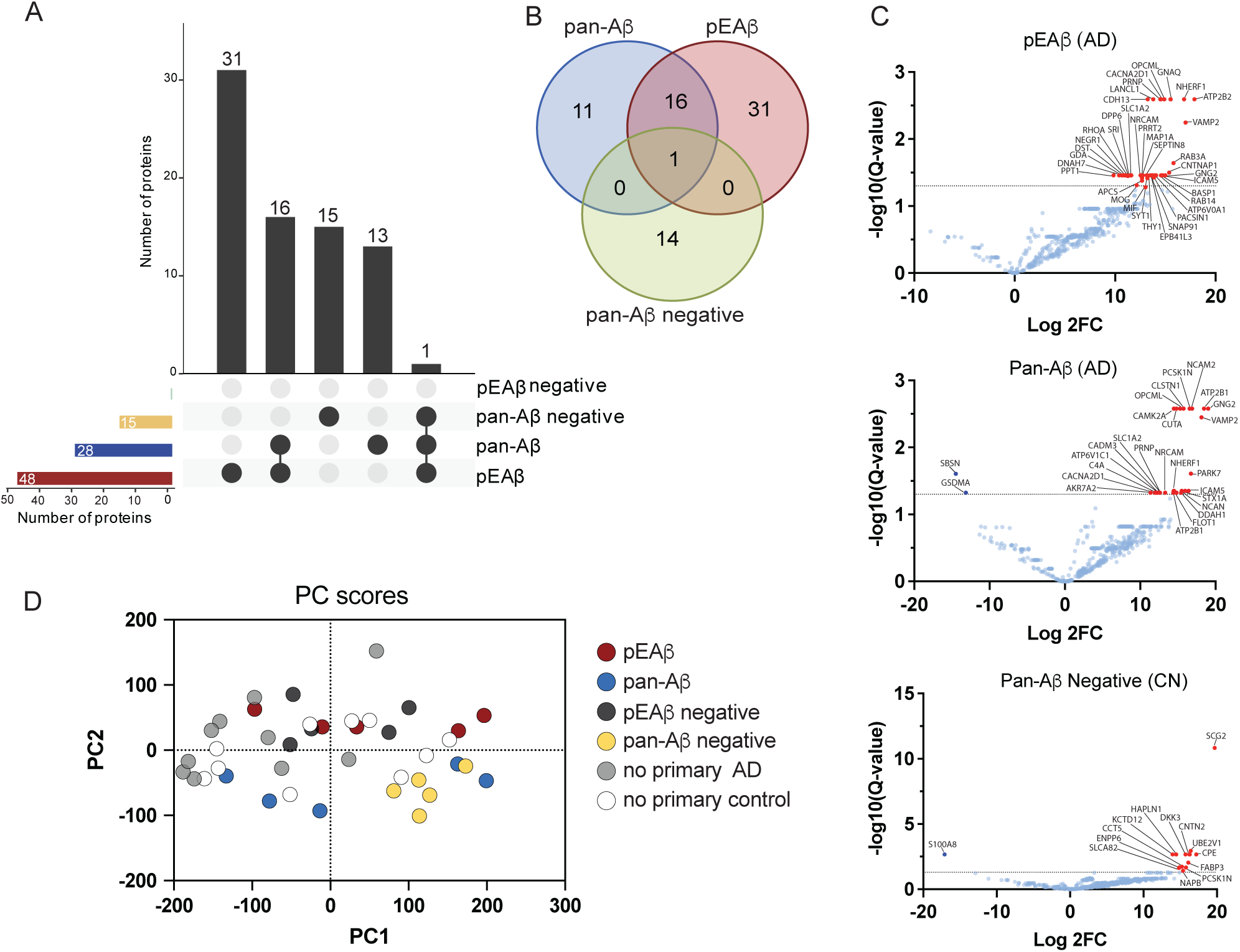
pEAβ and pan-Aβ proximity labeled proteins in human hippocampus. (A) Upset plot showing significantly abundant proteins across all experimental groups, with shared and unique proteins indicated on the plot. A total of 48 proteins were identified in the pEAβ group, including 31 unique proteins, 16 shared with the pan-Aβ group, and one shared among the pEAβ, pan-Aβ, and pan-Aβ–negative groups. The pan-Aβ group contained 28 identified proteins, of which 13 were unique. No proteins were detected in the pEAβ–negative group, while 16 were identified in the pan-Aβ–negative group, with 15 unique to that condition. (B) Venn diagram depicting unique and overlapping enriched proteins among BAR conditions. Unique proteins were identified for pEAβ (31), pan-Aβ (11), and pan-Aβ–negative (14) groups. Overlapping proteins were observed between the pan-Aβ and pEAβ groups (16), and among all three groups (1). Together, these findings suggest that pEAβ and pan-Aβ engage in overlapping core interactomes while also recruiting species- and disease-specific binding partners. (C) Volcano plots showing differential protein abundance between capture conditions, expressed as log₂ fold change (log₂FC) versus –log₁₀(Q-value). Each point represents a quantified protein, with significantly enriched proteins highlighted in red. (D) Principal component analysis (PCA) plot of proteomic profiles across all samples, with each point representing a biological specimen. Proximity along the axes indicates similar expression patterns, while separation reflects distinct proteomic signatures. The first two principal components explain 37% (PC1) and 7.8% (PC2) of total variance. Separation along PC1 demonstrates that distinct Aβ species drive divergent proteomic profiles in the human hippocampus.

**Table 3.**
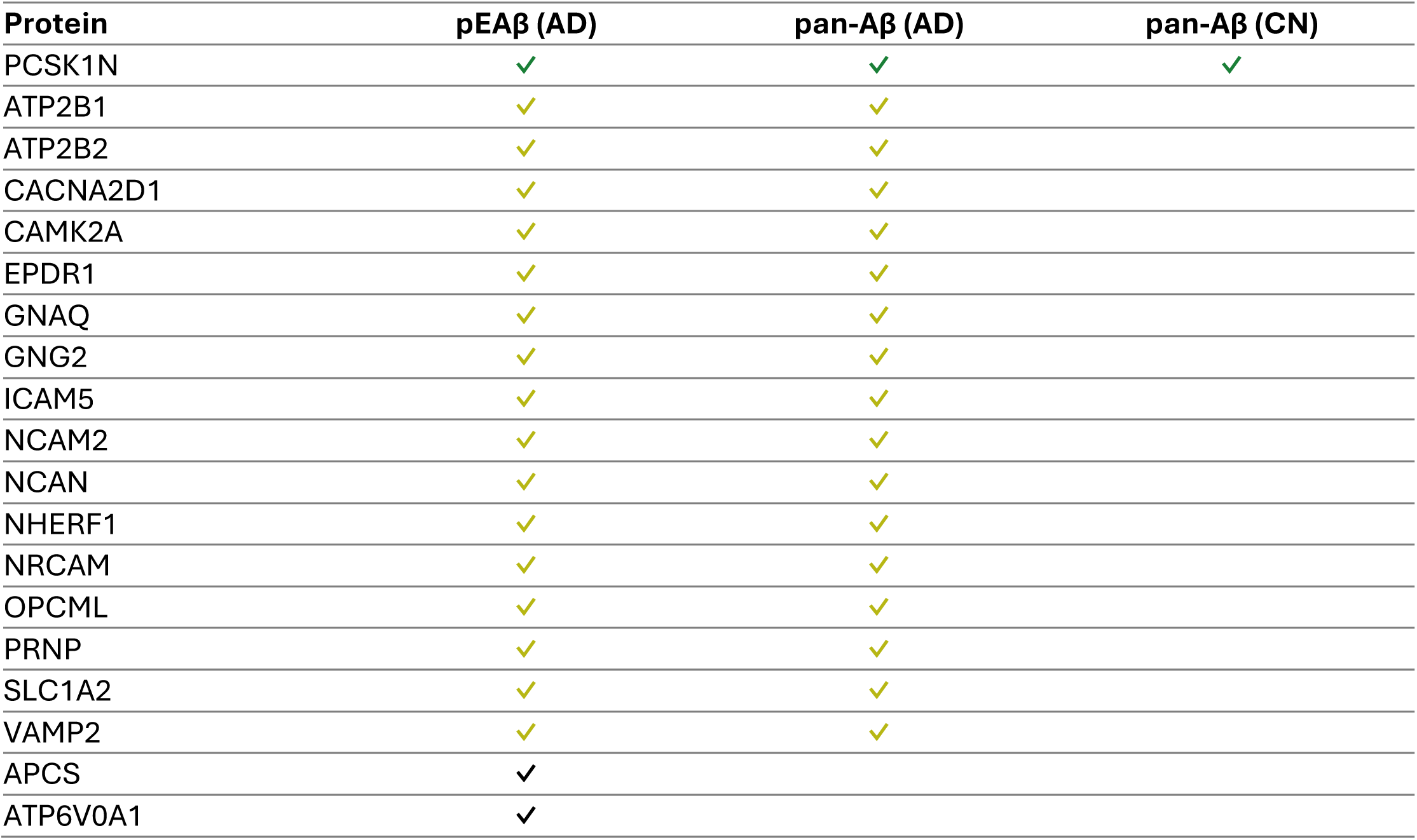
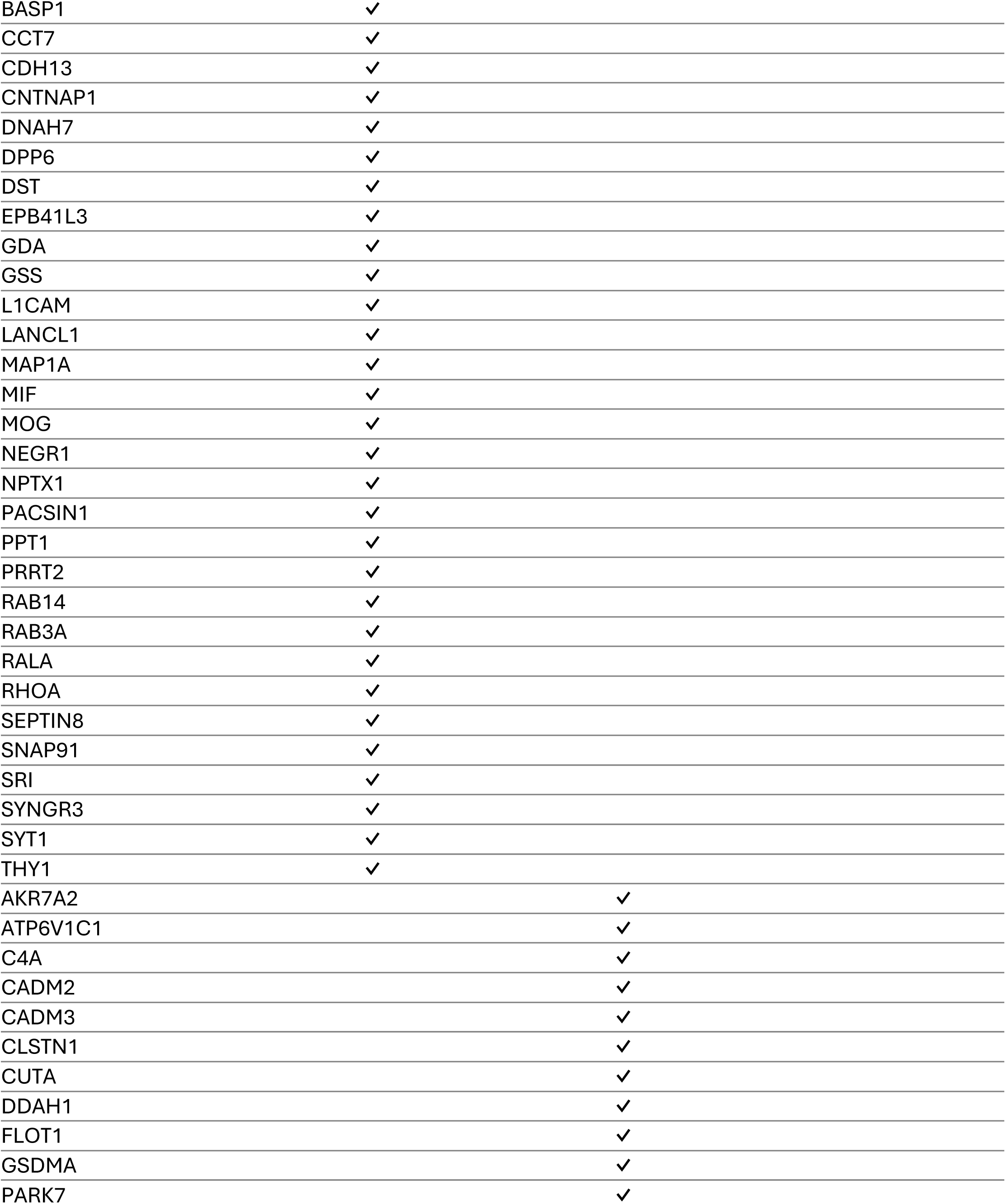
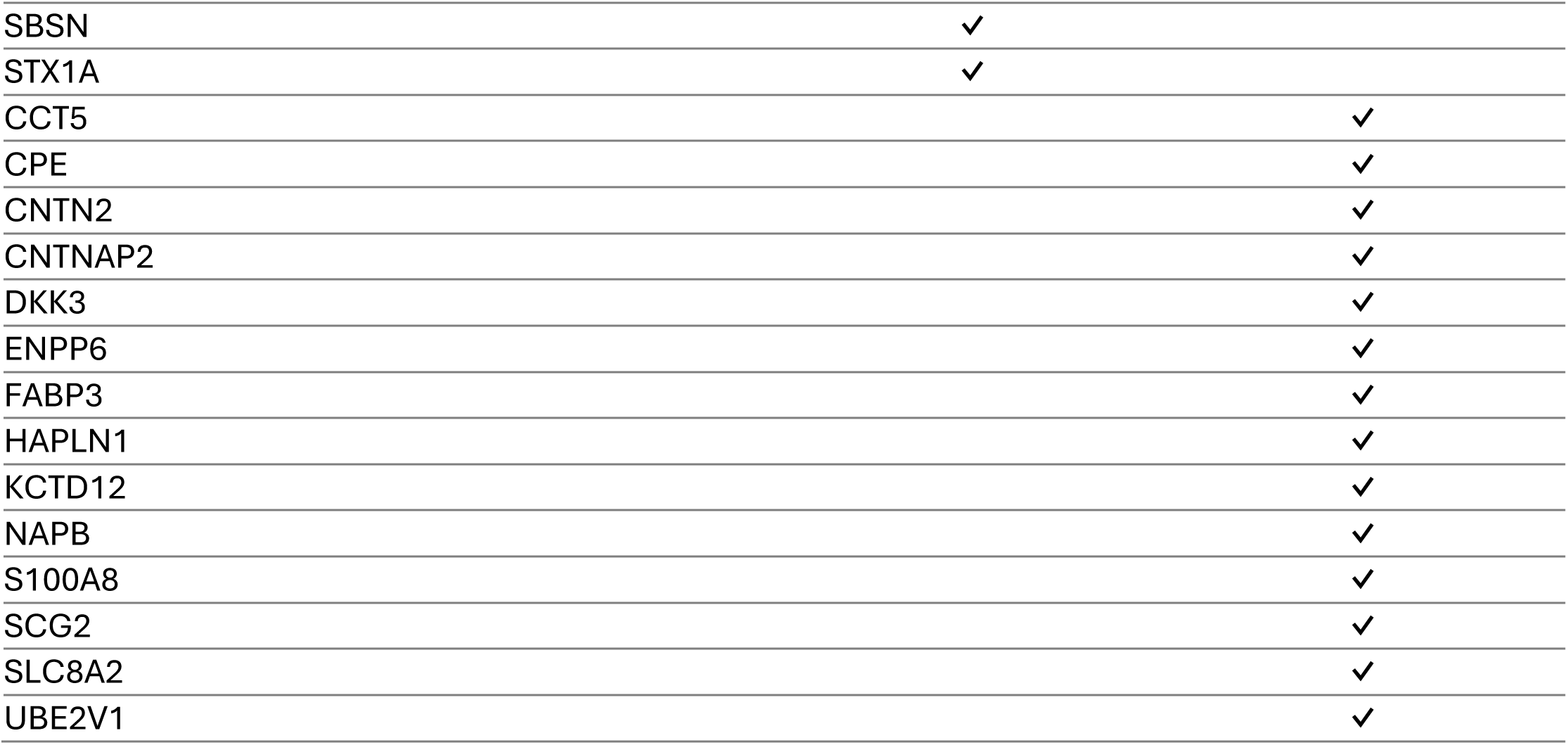
Presence/absence of proteins across Aβ capture conditions in human hippocampal tissue. Summary of proteins significantly enriched across Aβ capture conditions: pyroglutamate-modified Aβ in AD (pEAβ (AD)), pan-Aβ in AD (pan-Aβ (AD)), and pan-Aβ in CN tissue (pan-Aβ (CN)). Check marks indicate proteins identified in the corresponding capture condition; blank cells indicate proteins not detected in that condition. Green check marks indicate proteins shared across all three conditions; yellow check marks indicate proteins shared between pEAβ (AD) and pan-Aβ (AD). Proteins unique to a single condition are in black. Proteins are not listed in any particular order and are presented for ease of visual comparison.

Similarly, shared interactors were also detected between groups, with 16 shared proteins among AD pEAβ and AD pan-Aβ capture groups and one single shared protein (PCSK1N) among AD pEAβ, AD pan-Aβ, and CN pan-Aβ capture groups. These data demonstrate that unmodified Aβ and pEAβ participate in overlapping core interactomes while also engaging variant-specific protein interactomes.

Hierarchical clustering of protein abundance patterns revealed several distinct protein signatures that distinguished the Aβ specific interactomes (Fig. 4A). One major cluster consisted of proteins highly enriched in the pEAβ AD capture, demonstrating a discrete cluster that was minimally detected in pan-Aβ AD or CN samples. A second prominent cluster contained proteins selectively enriched in pan-Aβ AD captures. A third protein cluster represented a shared interactome, comprising proteins that displayed high abundance in both AD pEAβ and AD pan-Aβ groups, suggesting common molecular pathways engaged by both Aβ species. Finally, a small cluster of proteins was enriched only in the CN pan-Aβ condition, representing low Aβ burden interactors. Together, these protein level clusters demonstrate that pEAβ and pan-Aβ engage partially overlapping but distinct protein networks.

**Figure 4.**
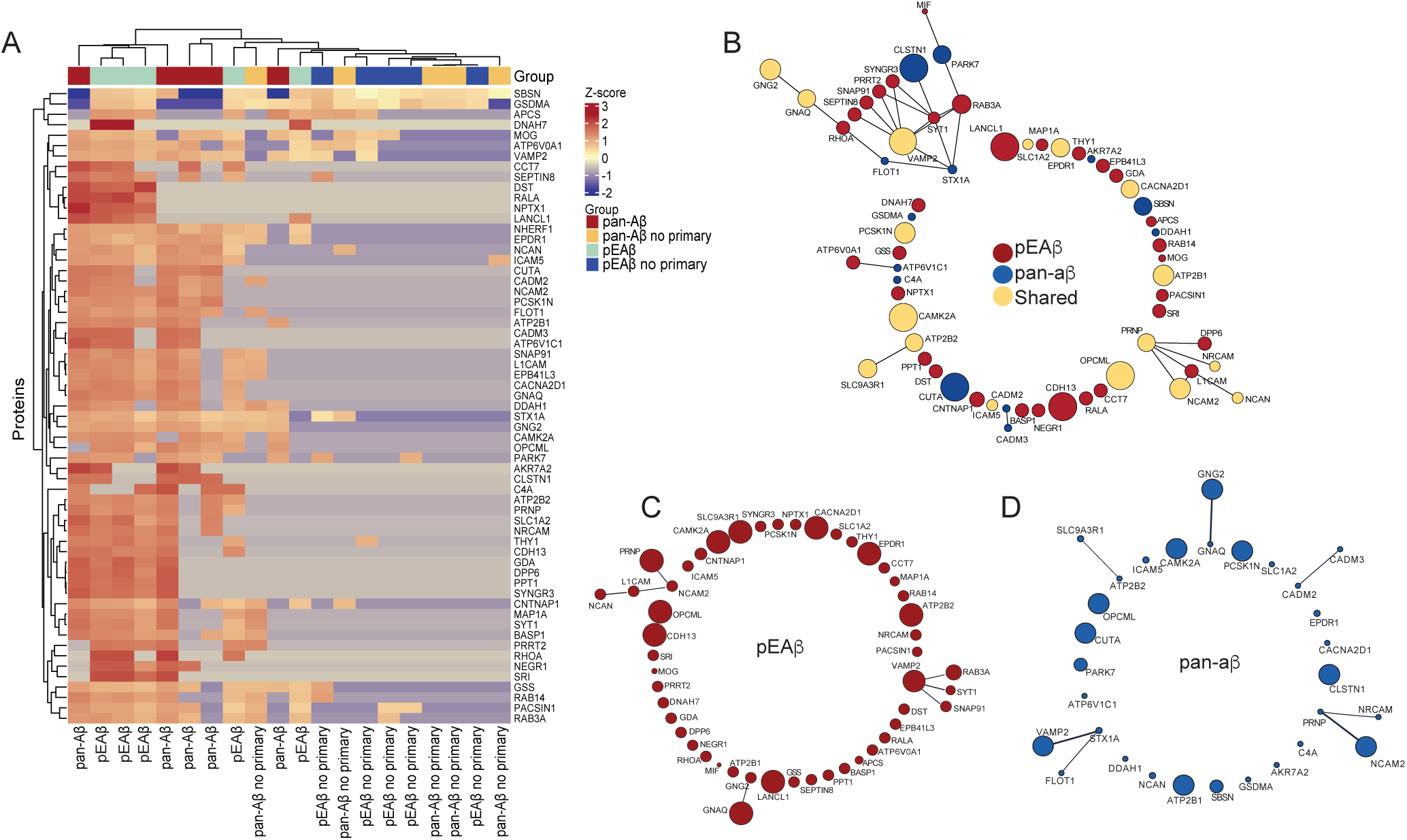
Analysis of pEAβ- and pan-Aβ–associated interacting proteomes in the human hippocampus. (A) Hierarchical clustering heatmap of Z-score–normalized protein abundance across pEAβ, pan-Aβ, and their respective no-primary antibody controls. Both pEAβ and pan-Aβ samples display elevated Z-scores and distinct clustering patterns, indicating enriched protein abundance, whereas control samples exhibit low signal intensity and cluster separately on the right. This segregation demonstrates specific protein enrichment associated with Aβ species and minimal non-specific background labeling. (B) STRING protein–protein interaction network of significantly enriched proteins identified in pEAβ (red), pan-Aβ (blue), and those shared between both (yellow). Two central shared nodes, *PRNP* and *VAMP2*, were identified, suggesting that pEAβ and pan-Aβ converge on synaptic signaling and membrane-associated pathways. These shared interactors highlight common mechanisms underlying amyloid-induced neurotoxicity and synaptic dysfunction. (C–D) STRING interaction networks of significantly enriched proteins unique to pEAβ (C, red) and pan-Aβ (D, blue). Proteins enriched in pEAβ are primarily involved in presynaptic vesicle turnover and docking, whereas those associated with pan-Aβ are linked to cell adhesion and synaptic vesicle fusion. Node size reflects statistical significance (Q-value, FDR-corrected p-value), with larger nodes indicating higher significance (Q < 0.05). Edges represent known or predicted protein–protein associations, delineating functional clusters within each interactome.

Among proteins selectively enriched in pEAβ AD, we identified a prominent cluster involved in synaptic vesicle trafficking and cell junction assembly, including RAB3A, RHOA, SYT1, ATP6V0A1, SNAP91, RAB14, PPT1, and PRRT2[45], [46], [47], [48], [49], [50], [51], [52], [53] (Fig. 4B-D). These proteins are functionally linked to synaptic vesicle docking, actin cytoskeleton regulation, and vesicle acidification, suggesting a possible role for pEAβ in disrupting presynaptic vesicle turnover and synaptic stability. In contrast, proteins enriched specifically in the pan-Aβ interactome included STX1A, CADM2, CADM3, CLSTN1, FLOT1, and PARK7, most of which are localized to the presynaptic membrane and are involved in cell adhesion, synaptic vesicle fusion, and oxidative stress response. Shared proteins between both Aβ variants in AD included VAMP2 and PRNP, which emerged as central nodes in the interactome network. Additional shared proteins such as CAMK2A, ATP2B2, NCAM2, and CACNA2D1 are predominantly localized to the plasma membrane and are known regulators of synaptic calcium signaling and plasticity[54], [55], [56], [57]. These findings suggest that both Aβ species converge on key synaptic signaling pathways, while also engaging unique protein networks that may contribute to their aggregation dynamics and toxicity.

In CN tissue, differential protein enrichment in pan-Aβ revealed a distinct subset of interactors that may reflect early or physiological Aβ engagement. Specifically, we identified proteins involved in synaptic vesicle regulation (SCG2, NAPB, RAP1B), adhesion and axon guidance (CNTNAP2, CNTN2, HAPLN1)[58], [59], [60], [61]. Additionally, proteins included regulators of protein homeostasis and modification (UBE2V1, CCT5, PCSK1N), lipid and metabolism-related proteins (ENPP6), and signaling modulators (DKK3)[62]. Collectively, these proteins represent a network that is functionally enriched for synaptic transmission, membrane adhesion, and intracellular trafficking, suggesting that in CN brains, pan-Aβ primarily associates with pathways important for maintaining synaptic integrity and neuronal communication, rather than visible pathological processes.

### KEGG subnetwork analysis reveals Aβ variant specific pathway modules

Analysis of protein subnetworks within each group revealed distinct KEGG pathway enrichments. In pEAβ AD samples, four physical clusters were identified, with proteins mapping to pathways including cell adhesion molecules (L1CAM, NRCAM, NCAM2), synaptic vesicle cycle (RAB3A, SYT1, VAMP2), chemokine signaling pathway (GNAQ, GNG2, RHOA), and RAS signaling pathway (GNG2 and RHOA). In contrast, pan-Aβ interactors in AD tissue formed five clusters, enriched in SNARE-mediated vesicular transport (STX1A and VAMP2), the synaptic vesicle cycle (STX1A, VAMP2), and insulin secretion (STX1A, VAMP2). For cognitively normal tissue, pan-Aβ upregulated proteins were enriched in pathways involving cell adhesion molecules (CNTN2, CNTNAP2). Notably, while the three groups engaged distinct plasma-membrane-associated proteins, they also shared overlapping synaptic proteins, highlighting overlapping effects on synaptic signaling despite differences in upstream interactions.

### Ingenuity Pathway Analysis reveals broader pathway disruption in pEAβ compared to unmodified Aβ in AD human hippocampus

To explore the functional relevance of proteins identified in proximity to Aβ variants, we performed Ingenuity Pathway Analysis (IPA) on the statistically significant interactors that were identified. This enabled the identification of key activated pathways by analyzing 48 proteins enriched in pEAβ AD and 28 proteins enriched with pan-Aβ Aβ. The results revealed that pEAβ significantly upregulated proteins were enriched in 35 biological pathways, whereas pan-Aβ significantly upregulated proteins were enriched only in 13 pathways, highlighting a broader range of biological processes impacted by pEAβ (Fig. 5A-B).

**Figure 5.**
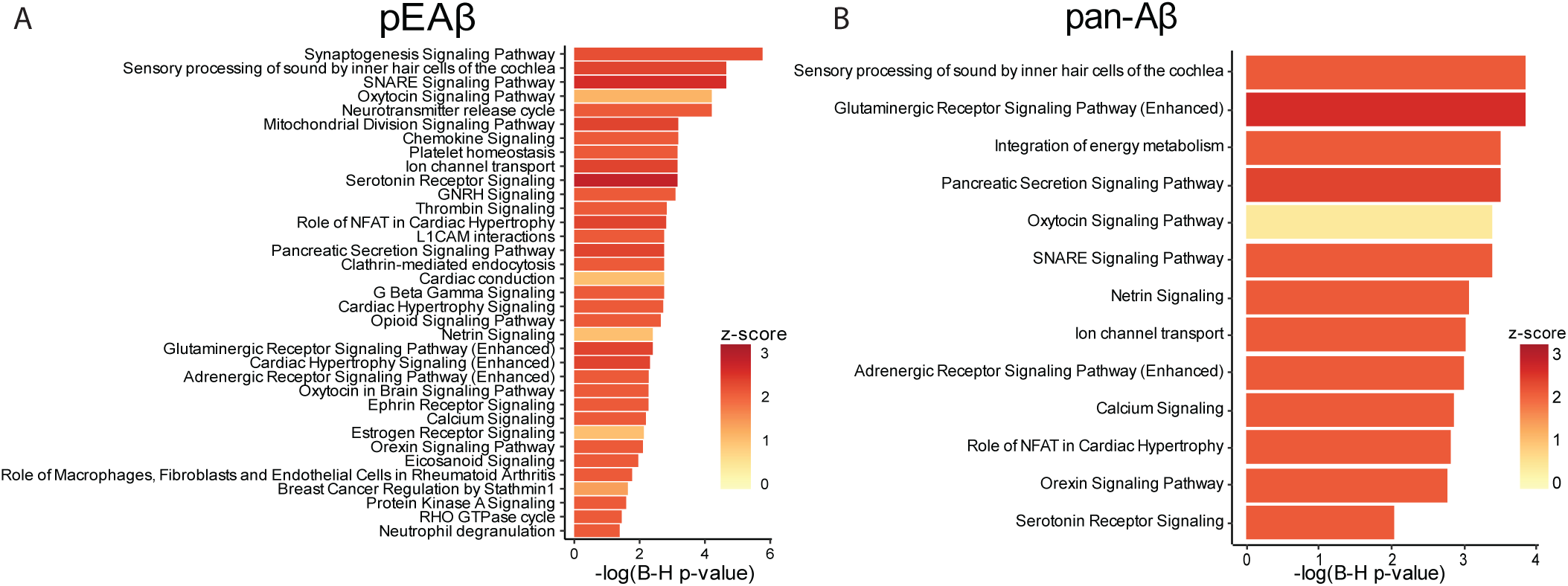
Ingenuity Pathway Analysis reveals activated pathways surrounding pEAβ and pan-Aβ fractions. (A) Bar plot demonstrating the top activated biological pathways associated with pEAβ in order of decreasing enrichment. A total of 35 pathways were significantly activated, including 22 unique to the pEAβ interactome. Prominent pEAβ-specific pathways include clathrin-mediated endocytosis, mitochondrial division signaling, and the neurotransmitter release cycle. (B) Bar plot of the top enriched biological pathways associated with pan-Aβ, arranged by decreasing enrichment. Thirteen pathways were significantly enriched in the pan-Aβ interactome. While no pathways were found to be unique to pan-Aβ, several were shared with pEAβ, including netrin signaling, SNARE signaling, and glutamatergic receptor signaling. The x-axis represents –log₁₀ (Benjamini–Hochberg adjusted *p*-values) for pathway enrichment, while the z-score denotes the predicted activation state, with values greater than 0 indicating pathway activation.

Among the most enriched pathways for pEAβ were the synaptogenesis signaling pathway (CACNA2D1,CAMK2A,CDH13,CNTNAP1,RAB3A,RALA,RHOA,SYT1,VAMP2), clathrin-mediated endocytosis (PACSIN1,SNAP91,SYT1,VAMP2), mitochondrial division signaling pathway (CACNA2D1,CAMK2A,GNAQ,RALA,RHOA), and neurotransmitter release cycle (RAB3A,SLC1A2,SYT1,VAMP2). These pathways suggest that pEAβ may disrupt key neuronal processes, including synapse formation, vesicle trafficking, and mitochondrial dynamics. Several pathways were shared between both Aβ variants, including the SNARE signaling pathway (CACNA2D1,CAMK2A,RAB3A,SNAP91,SYT1,VAMP2), glutaminergic receptor signaling pathway (CACNA2D1,CAMK2A,GNAQ,SLC1A2,VAMP2), and netrin signaling (CACNA2D1,CAMK2A,RHOA,VAMP2). Notably, no unique pathways were enriched in pan-Aβ that were not observed in pEAβ, reinforcing the idea that pEAβ engages a distinct and broader interactome. These findings emphasize the variant-specific biological roles of Aβ species in AD pathology and underscore the potential of pEAβ as a more disruptive molecular player.

## Discussion

Amyloid pathology in the human brain is highly heterogeneous and encompasses a spectrum of structural, biochemical, and molecular features[63]. At the ultrastructural level, Aβ assemblies can exist as soluble oligomers, protofibrils, or insoluble fibrils, which can exhibit distinct biological activities and neuropathological potential[63]. In addition to differences in aggregation state, Aβ peptides vary in length and undergo post-translational modifications, such as N-terminal truncation and pyroglutamate formation, that influence their stability, aggregation kinetics, and resistance to degradation[7]. Importantly, amyloid plaques are not composed solely of Aβ peptides but represent complex molecular assemblies that incorporate proteins, lipids, and extracellular matrix components [13]. These non-Aβ constituents are thought to modulate plaque maturation, persistence, and pathogenicity, underscoring the need to define the broader molecular environment surrounding distinct Aβ species within human brain tissue[13]. Therefore, understanding how specific Aβ variants engage surrounding proteins is critical for elucidating mechanisms underlying plaque formation, stabilization, and clearance. To address this, we used BAR, a proximity labeling technique that enables in situ capture of proteins located near a defined antigen within fixed tissue. This approach is particularly suitable for human postmortem samples, which are commonly formalin-fixed, and allows for the preservation of the native tissue environment while selectively labeling proteins within the immediate proximity of Aβ peptides. Using this spatially resolved strategy, we investigated the protein interactomes associated with unmodified Aβ and pEAβ in the human hippocampus. Consistent with our expectations, BAR revealed that distinct Aβ variants engage both shared and unique sets of interacting proteins. These findings support the idea that while different Aβ species participate in overlapping core molecular pathways, they also recruit variant-specific protein environments that may differentially contribute to amyloid aggregation, plaque maturation, and disease progression. We identified protein interactors of overlapping and divergent pathways that converge on synaptic and membrane-associated proteins.

Amyloid pathology in the human brain is characterized not only by the accumulation of Aβ peptides, but also by the complex molecular environments that surround and shape plaque formation. Both pan-Aβ and pEAβ were enriched in proteins associated with synaptic vesicle dynamics, consistent with the presynaptic origin of Aβ secretion [2]. However, these species vary in their biochemical properties and downstream consequences. Soluble monomeric pan-Aβ can be internalized, recycled, and degraded by Aβ-clearing mechanisms, whereas pEAβ is insoluble and aggregation-prone[2], [7], [64]. This insolubility may enable pEAβ to persist at synaptic membranes, promote further oligomerization, and bind to synaptic receptors[22]. Furthermore, we hypothesize that this process contributes to dysregulation of calcium homeostasis and triggers excitotoxic cascades [65]. In doing so, pEAβ impairs synaptic plasticity, disrupts long-term potentiation (LTP), and interferes with neurotransmitter release, consistent with the enrichment of synaptic vesicle cycle proteins (SYT1, VAMP2, RAB3A) and plasma membrane proteins identified in our dataset [47], [66].

Beyond synaptic vesicle machinery, our data indicate that distinct Aβ species are associated with different plasma membrane-related proteins that may converge on synaptic dysfunction. Distinct sets of plasma membrane proteins were identified across capture conditions, suggesting that pan-Aβ and pEAβ interact with different upstream partners. Notably, pEAβ− associated interactors included proteins linked to protein kinase A (PKA) signaling, a pathway known to regulate both synaptic transmission and tau phosphorylation[67], [68]. Although our data do not demonstrate direct pathway activation, these findings suggest pEAβ may be localized within membrane-associated protein networks relevant to synaptic dysfunction and tau-related processes. In parallel, the identification of adhesion molecules (L1CAM, NRCAM, NCAM2, CNTN2, CNTNAP2) indicates that both Aβ species are associated with proteins involved in synaptic connectivity and membrane integrity, consistent with prior reports linking Aβ to altered cell adhesion and synaptic maintenance[47], [56], [69].

In addition to synaptic and membrane-associated pathways, endocytic trafficking emerged as a key biological process associated with pEAβ pathology. One prominent pathway involved clathrin-mediated endocytosis (CME). CME is a fundamental pathway for internalizing membrane proteins and extracellular molecules and is strongly implicated in AD, particularly in relation to Aβ metabolism[70]. Since the QC activity that converts Aβ into pEAβ occurs within endocytic compartments, CME provides a critical trafficking route for Aβ to undergo this modification[71], [72]. Thus, impaired endosomal trafficking may not only disrupt recycling and synaptic vesicle turnover but also enhance the pathological conversion of full-length Aβ into pEAβ, further contributing to AD pathogenesis. This convergence highlights how defects in intracellular trafficking may amplify Aβ toxicity and promote plaque maturation.

Our pathway analysis also revealed upregulation of mitochondrial division signaling driven by pEAβ peptides, which may reflect compensatory responses to heightened energy demand or direct Aβ toxicity at mitochondria. Mitochondrial fragmentation is a well-described feature of Aβ and is closely tied to synaptic energy deficits[66], [73]. Here, excessive fission compromises neuronal energetics and heightens vulnerability to oxidative stress. The presence of chemokine signaling proteins (GNAQ, GNG2, RHOA) in both pan-Aβ and pEAβ protein networks further suggests that Aβ species influence inflammatory signaling. This finding aligns with prior evidence that Aβ can recruit microglia and astrocytes to amplify neuroinflammation[74]. These results emphasize that pEAβ is positioned between synaptic, mitochondrial, and inflammatory dysfunction.

PCSK1N was the only protein detected across all three capture conditions (pEAβ AD, pan-Aβ AD, and pan-Aβ CN), making it a particularly central node that may reflect a core feature of Aβ biology.

PCSK1N (proSAAS) is a secretory granule-associated protein that is predominantly expressed in the brain and neuroendocrine tissues [75], [76]. proSAAS demonstrates significant colocalization with Aβ plaques in human AD brain tissues, with strong localization observed mainly in dense core pathology and also evidence of accumulation in early-stage diffuse plaques, supporting an association across multiple stages of plaque maturation[75], [76]. Importantly, because BAR reports spatial proximity rather than direct binding, the consistent detection of PCSK1N across pEAβ AD, pan-Aβ AD, and pan-Aβ CN suggests that proSAAS may be a conserved component of the Aβ-adjacent microenvironment rather than strictly a disease-specific interactor. In our dataset, proSAAS showed the most significant enrichment in the pEAβ AD capture condition, suggesting that proSAAS may be preferentially concentrated within pEAβ-rich plaques, which are typically more compact and core-associated. This enrichment pattern is compatible with two reported interpretations: that proSAAS is a neuronal secretory/proteostasis player that may act locally to limit aggregation, and the interpretation that proSAAS displays increased retention within dense-core deposits; however, it may not necessarily reflect a functional interaction with Aβ. Together, these findings position PCSK1N as a shared “bridge” protein across physiological and pathological Aβ contexts, while its relative enrichment in pEAβ captures may track plaque maturation and the distinct microenvironment of pEAβ dominant pathology. A critical distinction emerging from this study is the dissimilarity between interactomes surrounding unmodified Aβ in cognitively normal and AD tissue. In cognitively normal tissue, we detected pan-Aβ interactors but observed no detectable pEAβ interactome. This finding suggests that early deposition engages synaptic and adhesion pathways without progressing to the more pathogenic pEAβ species. Thus, pEAβ associated protein networks may represent a later disease-specific stage that distinguishes normal physiological Aβ turnover from pathological Aβ persistence. This distinction underscores why pEAβ is an especially relevant therapeutic target in AD progression.

Cognitively normal samples revealed distinct interactomes depending on the Aβ species. Here, cognitively normal pan-Aβ proteins are related to cell adhesion molecules, suggesting a physiological role in maintaining synaptic contacts and neuronal membrane organization. Whereas cognitively normal pEAβ samples lacked detectable interactors, consistent with the absence of pEAβ deposition in healthy brains. This supports the concept that unmodified Aβ may serve homeostatic roles, while post-translational modifications to Aβ peptides demonstrate pathogenicity through altered aggregation propensity and expanded interaction networks[2], [77].

Several limitations exist in this study. First, the lack of statistically significant enrichment of major plaque-associated proteins, including amyloid precursor protein (APP), apolipoprotein E (ApoE), and clusterin (CLU). The 6E10 antibody used for the study does not recognize a purely Aβ-specific target, as research has shown that 6E10 immunoreactivity may represent a mixture of deposited Aβ, APP-derived fragments, and other APP-containing material, rather than exclusive insoluble Aβ deposits [78], [79]. Here, the BAR proximity-based method used is not a direct binding assay. Because the technique deposits biotin onto proteins adjacent to the antibody-bound antigen, the proteins recovered by mass spectrometry reflect the local environment surrounding the 6E10-positive signal rather than only proteins that directly bind Aβ. In this context, if 6E10 recognizes APP-related material in addition to insoluble Aβ, the resulting proteome may shift toward a broader APP/Aβ proximal proteome and thereby dilute other interactions specific to Aβ only, thus APP may be a part of the antibody-recognized antigenic environment. Concerning other major plaque-associated proteins such as ApoE and clusterin, ApoE is known to regulate extracellular Aβ metabolism, clearance, and deposition, while clusterin functions as an extracellular chaperone that interacts with aggregation-prone amyloid species[80], [81]. However, these interactions may be transient, conformation-dependent, or enriched in soluble extracellular complexes rather than stably represented within the immediate environment of 6E10-direct labeling at the time of fixation. Therefore, the absence of significant enrichment in this study should not be interpreted as evidence that these proteins are unimportant in Aβ biology, but rather that they were not robustly captured under the conditions used here. An additional limitation of this study is the relatively small sample size (n=10), which may have reduced statistical power and limited the ability to detect proteins with modest enrichment. Although the cohort was sufficient to distinguish differences between Aβ species, the specific proteins identified or reaching statistical significance could differ in a larger study. In addition, cases were grouped primarily by disease status, and the present study was not designed or powered to evaluate independent effects of other potentially important biological variables such as age, sex, ethnicity, or regional brain differences. Therefore, the extent to which these variables influenced the observed results could not be determined in the current cohort. A further limitation of this study is that the analysis was restricted to the hippocampus. Although this region is highly relevant to AD and offers a consistent anatomical framework for comparison, Aβ deposition arises earlier in neocortical regions during disease progression[82]. Therefore, the interactomes described here may not fully represent the earliest molecular changes associated with Aβ deposition and instead may reflect hippocampal or later-stage plaque-associated protein environments. Furthermore, CAA was observed in all AD cases, and the BAR approach and antibody selection used in this study do not distinguish vascular Aβ from plaque-associated Aβ. As a result, the interactomes identified here may reflect a combination of plaque-associated and vascular amyloid environments.

Taken together, these findings carry several therapeutic implications. While current immunotherapies such as Donanemab directly target pEAβ, our data highlight additional intervention points. These include pathways regulating endocytic trafficking, synaptic vesicle cycling, kinase signaling (including PKA), mitochondrial dynamics, and inflammatory responses. Targeting these downstream processes may complement existing antibody-based strategies and mitigate broader network-level consequences of pEAβ accumulation. By distinguishing pan-Aβ and pEAβ interactomes in human hippocampal tissue, this study provides a molecular framework that accounts for the biochemical heterogeneity of Aβ variants and their distinct pathological roles.

## Conclusion

In conclusion, our results suggest that pEAβ functions not only as a toxic peptide but also as a hub for dysregulated protein networks at synapses, plasma membranes, and intracellular trafficking compartments. By mapping distinct and overlapping interactomes, we demonstrate that pan-Aβ participates in physiological and early pathological processes, whereas pEAβ engages a set of pathways that amplify synaptic failure, tau phosphorylation, mitochondrial stress, and neuroinflammation. Importantly, no significant interactors were detected in the pEAβ capture condition in CN hippocampus, consistent with minimal pEAβ deposition in healthy tissue and supporting the interpretation that pEAβ-associated networks represent a disease-linked, pathological stage of Aβ. In contrast, comparison of unmodified Aβ profiles in AD and CN tissue revealed distinct networks, revealing proteomic profile shifts of baseline molecular pathways that become altered in the diseased state. This work advances our understanding of why pEAβ is a compelling therapeutic target and demonstrates how spatial interactome profiling can uncover network-level vulnerabilities in AD.

## Supporting information

Additional file 1 dot blot

Additional file 2 specimens_manifest_diff_PCA_IPA

## List of abbreviations

Aβ: Amyloid beta
Aβ40: Amyloid beta 1–40
Aβ42: Amyloid beta 1–42
ACE: Angiotensin-converting enzyme
ACN: Acetonitrile
AD: Alzheimer’s disease
AmBic: Ammonium bicarbonate
ApoE: Apolipoprotein E
APP: Amyloid precursor protein
ARIA: Amyloid-related imaging abnormalities
AZSAND: Arizona Study of Aging and Neurodegenerative Disease
BAR: Biotinylation by antibody recognition
BBB: Blood-brain barrier
CAA: Cerebral amyloid angiopathy
Clu: Clusterin
CME: Clathrin-mediated endocytosis
CN: Cognitively normal
DAB: 3,3′-diaminobenzidine
DDA: Data-dependent acquisition
DM: Dilution media
DTT: Dithiothreitol
ECE-1: Endothelin-converting enzyme-1
FDA: Food and Drug Administration
GLM: General linear model
HCD: Higher-energy collisional dissociation
HRP: Horseradish peroxidase
IAA: Iodoacetamide
IDE: Insulin-degrading enzyme
IHC: Immunohistochemistry
NEP: Neprilysin
pE: Pyroglutamate
pEAβ: Pyroglutamate-modified amyloid beta
PGP: Pyroglutamyl peptidase
PMI: Post-mortem interval
QC: Quality control
QPCT: Glutaminyl-peptide cyclotransferase
TIC: Total ion current

## Declarations

### Ethics approval and consent to participate

All procedures involving human tissues were approved by the Institutional Review Board (IRB) of Banner Sun Health Research Institute brain bank for the Arizona Study of Aging and Neurodegenerative Disease as a part of the Brain and Body Donation Program (20120821: 08/01/2025).

### Consent for publication

The study subjects provided informed consent according to the Banner Sun Health Research Institute brain bank from the Arizona Study of Aging and Neurodegenerative Disease IRB protocol.

### Availability of data and material

All data supporting the findings of this study are available within the paper and supplementary information. The datasets generated and analyzed during the current study are available in the Proteomics Identification Database (PRIDE) repository.

### Competing interests

Alia O. Alia, Kristy Urquhart, Hannah Carson, Bryan Killinger, Christopher Janson and Liudmila Romanova declared no competing interests or conflicts.

### Funding

The study was funded by Searle Innovator Grant by Searle Foundation to Dr. Liudmila Romanova.

### Authors contributions

Conceptualization, LR, AA, BK, CJ. Methodology, validation, data analysis, and investigation, AA, LR, BK, KU, HC. Interpretation, AA, LP BK, CJ, KU. Writing original draft, AA. Reviewing and editing, AA, LR, BK, CJ. Supervision, LR, BK. Funding acquisition, LR, BK. Visualization, AA, LR, BK.

## Acknowledgements

We are grateful to the Banner Sun Health Research Institute Brain and Body Donation Program of Sun City, Arizona, for the provision of human biological materials. Proteomics services were performed by the Northwestern Proteomics Core Facility, supported by NCI CCSG P30 CA060553 awarded to the Robert H Lurie Comprehensive Cancer Center, instrumentation award (S10OD025194) from NIH Office of the Director, and the National Resource for Translational and Developmental Proteomics supported by P41 GM108569. Data processing and analysis support was performed by the University of Illinois at Chicago Research Informatics Core (UIC RIC).

## Additional Files

1. Biotin Dot blots and antibody specificity dot blots
2. Table of all proteins identified and significant proteins surrounding each Aβ variant, including localization within the cell, molecular function, and biological process. IPA pathway datasets and PCA coordinates.

## Notes

### Competing Interest Statement

The authors have declared no competing interest.

