## Additional file 1 dot blot for "Proximity labeling reveals unique and shared interactomes of unmodified and pyroglutamate amyloid beta in human hippocampus in Alzheimer’s disease": Additional file 1_Dot_blots.docx

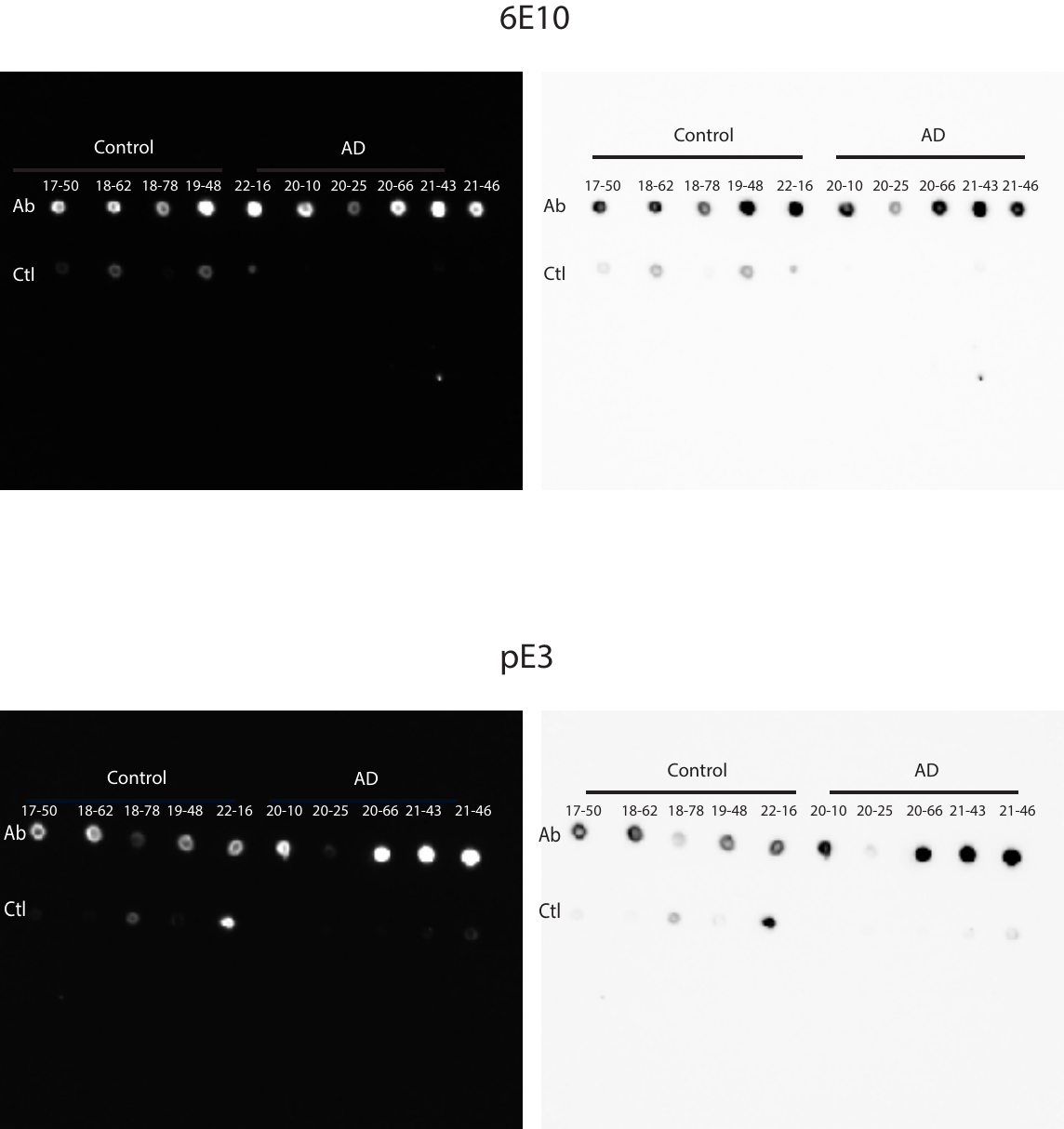

**Figure 1A. Biotin Dot blot** to estimate target enrichment after performing BAR on human hippocampus tissue (AD and Control). PVDF membranes were spotted with 1 µL of each BAR sample and biotinylated material was detected using an avidin–biotin complex (ABC) followed by ECL development. Top panel: representative dot blot of biotinylated material from specimens labeled with 6E10 antibody against unmodified amyloid beta. Bottom panel: representative dot blot of biotinylated material from the pE3 peptide antibody against pyroglutamate amyloid beta.

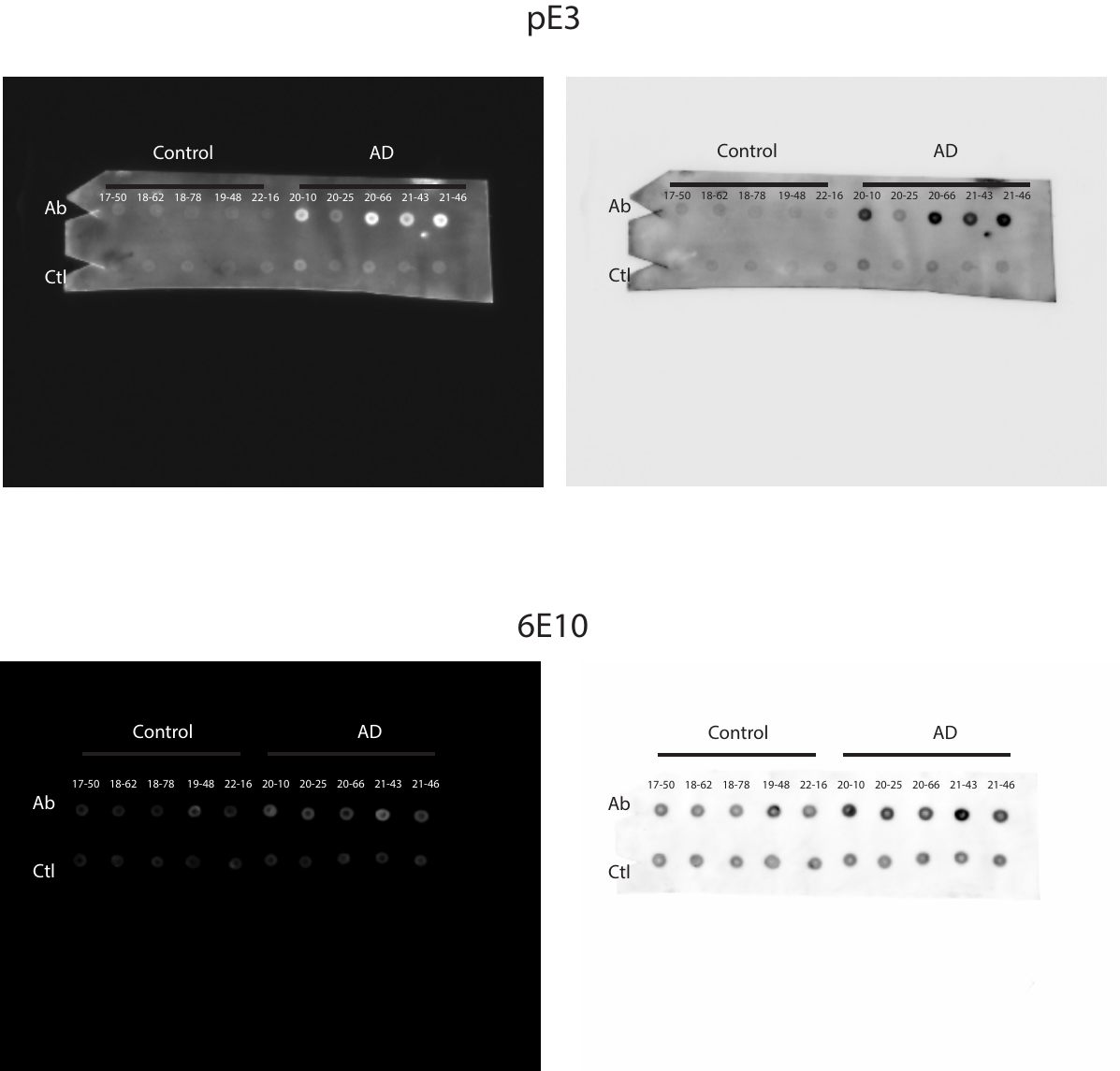

**Figure 2A. Antibody detection dot blots** to estimate antibody (antigen) enrichment after performing BAR on human hippocampus tissue (AD and Control). PVDF membranes were spotted with 1 µL of each BAR eluate and probed with the corresponding primary antibody (pE3 peptide or 6E10) followed by HRP-conjugated secondary antibody and ECL detection. Top panel: representative dot blot of eluates from pE3 peptide labeling. Bottom panel: representative dot blot of eluates from 6E10 labeling.
